# Fucose is an essential feature in cryoprotective polysaccharides

**DOI:** 10.1101/2023.10.13.562212

**Authors:** B. M. Guerreiro, P. Concórdio-Reis, H. Pericão, F. Martins, X. Moppert, J. Guézennec, J.C. Lima, J.C. Silva, F. Freitas

## Abstract

Biological cryopreservation often involves using a cryoprotective agent (CPA) to mitigate lethal physical stressors cells endure during freezing and thawing, but effective CPA concentrations are cytotoxic. Hence, natural polysaccharides have been studied as biocompatible alternatives. Our current investigation studied 26 natural polysaccharides as potential CPA, probing correlations between post-thaw metabolic viability (PTV) of cryopreserved Vero cells and monomeric composition. The best performing cryoprotective polysaccharides contained significant fucose amounts, resulting in average PTV 2.8-fold (up to 3.1-fold) compared to 0.8-fold and 2.2-fold for all non-cryoprotective and cryoprotective polysaccharides, respectively, outperforming the optimized commercial CryoStor™ CS5 formulation (2.6-fold). Stoichiometrically, a balance between fucose (18–35.7 mol%), uronic acids (UA) (13.5–26 mol%) and high molecular weight (MW > 1 MDa) generated optimal PTV. To deconvolute multiple variable effects, principal component analysis (PCA) coupled to *K*-means clustering was performed. Two major mechanisms of action explained PTV variability: a charge-dependent effect of contrasting charged uronic acid and neutral monomer compositions, and a MW-scaled charge-independent mechanism exclusively attributed to fucose. Ultimately, our research showed the critical role neutral fucose plays in enhancing cellular cryopreservation outcomes, disputing previous assumptions of polyanionicity being the sole governing predictor of cryoprotection, highlighting the potential of fucose-rich polyanionic polysaccharides.

## 1. Introduction

Cryopreservation, the process of freezing cells, tissues and organs to preserve them for future use, has revolutionized many fields of science (Bojic et al., 2021) and medicine (Giwa et al., 2017). During the freezing of biological material, the thermodynamics governing a phase transition will promote ice nucleation and growth, which are lethal physical stressors if a cryoprotective agent (CPA) is not used. Both small-molecule and polymeric CPAs are molecules with the ability to disrupt crystal formation by establishing strong interactions with water molecules (Murray & Gibson, 2022). This disruption can occur by freezing avoidance mechanisms, such as ice growth inhibition, ice recrystallization inhibition (IRI), dynamic ice shaping and thermal hysteresis; or at the nucleation level, where ice nucleation promotion by proteins (INPs) (Qiu et al., 2019; Roeters et al., 2021) and polysaccharides (Guerreiro et al., 2022) has been observed as a strategy of freezing tolerance. In addition to ice damage, the phase transition will generate aggressive osmotic flows of water and solutes, forcing cells to undergo strenuous volumetric changes that can lead to cell lysis (Mazur, 2010). This is usually controlled by osmotic regulators (permeant or impermeant CPAs) that counteract the differences in tonicity between the cell membrane. While cryopreservation has been widely used for decades, the issue of CPA cytotoxicity is as relevant as ever. Gold standard cryoprotectants such as DMSO and glycerol only show an appreciable cryoprotective effect at inherently cytotoxic concentrations. This results in overly complex cryopreservation protocols (Benson et al., 2018) that require several CPA loading-unloading cycles and washout steps (Guan et al., 2012).

To address these challenges, new cryoprotective alternatives have been explored. Among several synthetic (Gilfanova et al., 2021; Stubbs et al., 2020) and natural polymers (Carillo et al., 2015; Guerreiro et al., 2020; Khudyakov et al., 2015; X. Sun et al., 2022), biologically produced antifreeze proteins (AFPs) have excellent cryoprotective performance (Yeh & Feeney, 1996) but their purification and isolation is not cost-effective. Conversely, bio-based polysaccharides produced by microorganisms can leverage a byproduct-based circular economy of added value production (Freitas et al., 2011; Matsumura et al., 2022) that can rival AFP implementation in cryoprotective formulations both in cost and performance (Guerreiro, Silva, Torres, et al., 2021).

In addition to their biocompatibility and cryoprotective potential, some natural polysaccharides possess a consortium of bioactive functions, namely osmoregulatory (Khudyakov et al., 2015; Liu et al., 2013), anti-inflammatory (Araújo et al., 2023; Carillo et al., 2011; Khudyakov et al., 2015), antitumoral (Jiang et al., 2006; Kazak Sarilmiser & Toksoy Oner, 2014), immunoregulatory (Carillo et al., 2011; Ruiz-Ruiz et al., 2011), wound healing (Concórdio-Reis et al., 2020; Cui et al., 2022), gel-forming (Akan & Oner, 2021; Araújo et al., 2023; Fialho et al., 2019), emulsifying (Baptista et al., 2022; X. Chen et al., 2019; Nakamura et al., 2004) and neuroprotective (Xu et al., 2022) effects, among others, that further support their broad use in the preservation of complex biological systems which benefit from increased multifunctionality.

For instance, the fucose-rich extracellular polysaccharide (EPS) FucoPol has so far demonstrated strong non-colligative ice growth disruption and re-shaping (Guerreiro et al., 2020), nucleation promotion and stochastic narrowing (Guerreiro et al., 2022), versatile use in cryoprotective formulas of variable composition and different cell lines (Guerreiro, Silva, Torres, et al., 2021), and an antioxidant effect (Guerreiro, Silva, Lima, et al., 2021) that counteracts the ROS-mediated, cryopreservation-induced mitochondrial permeability transition pore (MPTP) opening that leads to delayed post-thaw cell death (Thomson et al., 2009). All these properties, associated with a high-MW, high viscosity and shear-thinning behavior (Torres et al., 2015) that facilitates nutrient diffusion and promotes cellular attachment by natural design (Limoli et al., 2015), conceive a molecule capable of maximizing the post-thaw viability of cryopreserved biologicals.

However, the structure-function relationships between polysaccharides and their ability to protect cells during cryopreservation is not yet fully understood (Casillo et al., 2017, 2021). Deconvoluting these relationships requires a comprehensive understanding of their physicochemical properties, such as MW, chemical structure, charge density, spatial conformation, and sequence patterns (Gray et al., 2019).

In this work, we explored the cryoprotective potential of several natural polysaccharides, handpicked to generate a spectrum of known compositions and molecular weights. The metabolic and histological viability of cryopreserved Vero cells was determined in freezing media containing different polysaccharides supplemented with 10% DMSO. It is widely accepted that polyanionicity plays a major role in most biologically active polysaccharides, including cryopreservation (Matsumura et al., 2022). The property is mostly established by the presence of uronic acid (UA) monomers that contain a deprotonated carboxylate moiety at physiological pH (H. M. Wang et al., 1991), but may also be conferred by sulfate, phosphate or other acyl modifiers in the polymer chain. However, some research studies have shown that fucose-containing polysaccharides show an appreciable effect towards cryopreserved cells (Guerreiro et al., 2020; Guerreiro, Silva, Torres, et al., 2021), and that cryoprotective molecules need not necessarily be charged to exert an influence (Rajan et al., 2016; Y. Sun et al., 2021), raising suspicion on possible governing mechanisms that lay under the umbrella label of cryoprotective action other than formal charge. Thus, we cross-correlated the compositional and structural features of each polysaccharide with its *in vitro* performance using principal component analysis (PCA) to draw insights of potential predictors of variability, with the intent of gaining insight towards being able to develop improved cryopreservation formulations that are safe, effective, and widely applicable, based on intentional design.

## Hypothesis

It is widely accepted that polyanionicity plays a major role in most biologically active polysaccharides, including cryopreservation (Matsumura et al., 2022). The property is mostly established by the presence of UA monomers that contain a deprotonated carboxylate moiety at physiological pH (H. M. Wang et al., 1991), but may also be conferred by sulfate, phosphate or other acyl modifiers in the polymer chain. However, some research studies have shown that fucose-containing polysaccharides show an appreciable effect towards cryopreserved cell survival (Guerreiro et al., 2020; Guerreiro, Silva, Torres, et al., 2021), and that cryoprotective molecules need not necessarily be charged to exert an influence (Rajan et al., 2016; Y. Sun et al., 2021). Here, we hypothesize that, from the possible governing mechanisms that lay under the umbrella label of cryoprotective action other than formal charge, neutral monomers in the chain, especially fucose, may play a bioactive role other than chain linker functionality.

## 2. Materials and Methods

### 2.1. Cell lines, media & reagents

Low-glucose Dulbecco’s Modified Eagle’s Medium (DMEM, #D5030) supplemented with 10% fetal bovine serum (FBS, #10270106 Invitrogen) and 1% penicillin-streptomycin (#15140122, Invitrogen) was used to proliferate Vero cells (monkey kidney epithelium, ATCC^®^ CCL-81^TM^). Cells were kept in incubation at 37°C, 5% CO_2_ and experiments initiated at 70-80% cell confluency. CryoStorT^M^ CS5 was obtained from BioLife Solutons (#18054).

### 2.2. Polysaccharides

M-rich alginate (sodium salt from brown algae, Sigma 9005-38-3), G-rich alginate (sodium salt from *Laminarea hyperborea*, rich in guluronic acid, BDH Chemicals 30105), carrageenan κ (Fluka 22048), carrageenan κ+λ (type I commercial grade, predominantly κ, Sigma 9000-07-1), dextran 40 (from *Leuconostoc spp.*, Sigma 9004-54-0), Ficoll^®^400 (GE Healthcare 17-0300-10), Ficoll^®^70 (GE Healthcare 17-0310-10), guar (Fluka 09999), hyaluronic acid (sodium salt, Hyasis 850P, Novozymes 4100004), xanthan (from *Xanthomonas campestris*, Sigma 11138-66-2), starch (Acrös Organics 9005-84-9), pectin (from citrus fruits, Sigma 9000-69-5) and agar (bacteriological, Scharlau 07-004-500) were obtained commercially. The biotechnologically produced polysaccharides were obtained from the microorganisms presented in Table 1.

**Table 1.**
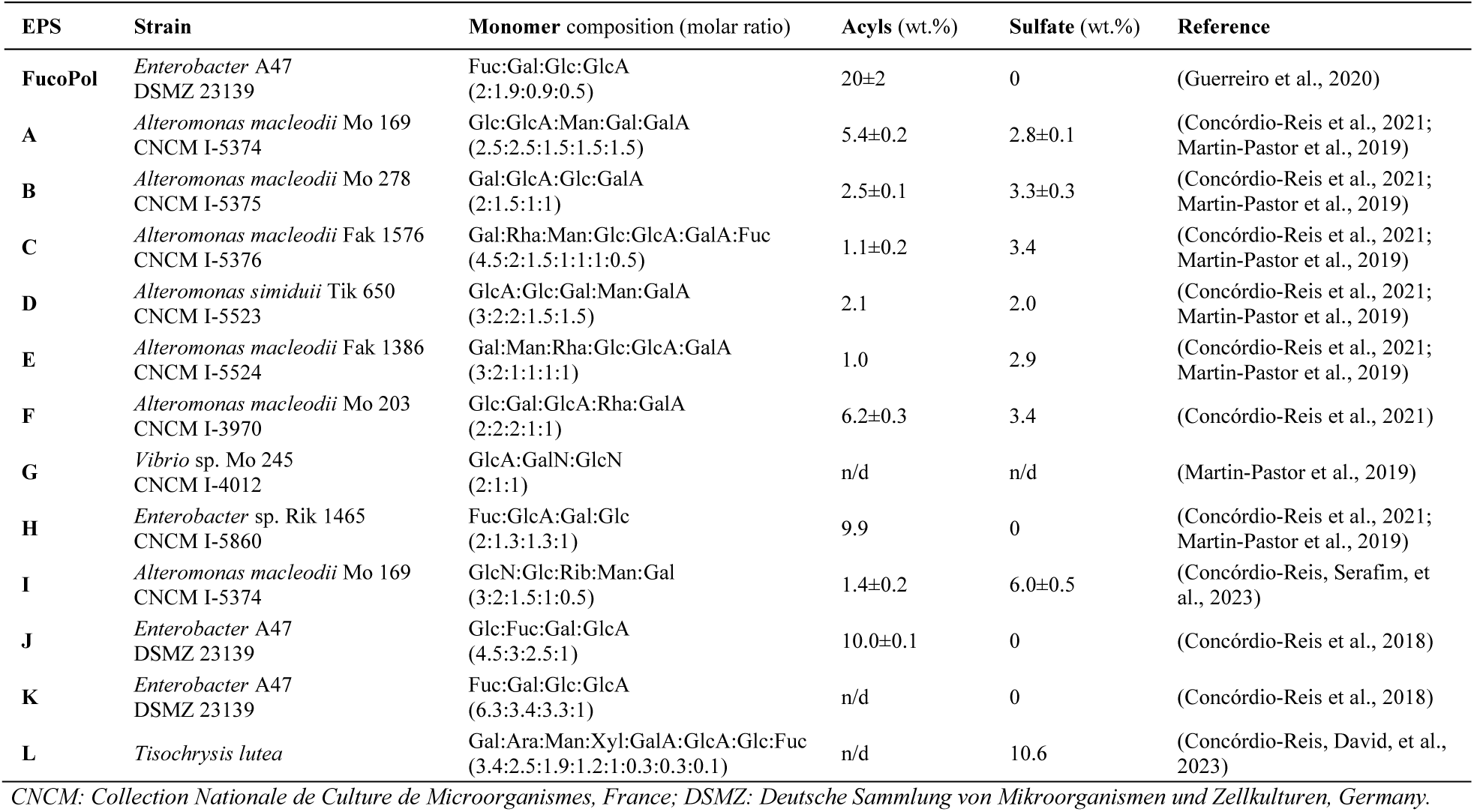
Biological origin of each polysaccharide used in the present study, and their monomer composition.

### 2.3. Cell monolayer cryopreservation

Vero cells were seeded in 96-well microplates with a cell density of 40,000 cells/well in low-glucose DMEM, and incubated at 37°C, 5% CO_2_ for 24 h. Then, the adhered cells were exposed to a freezing medium containing: 10% v/v DMSO and each test polysaccharide at either 0.5, 0.25 or 0.125 wt.% dissolved in DMEM, for a total volume of 100 μl in each well. Each condition was replicated four times. The addition of DMSO is performed in the solution preparation phase rather than direct microwell injection to avoid in-well DMSO heterogeneity and minimize pre-freeze exposure time. The strategic supplementation of 10% DMSO was designed to compensate for the shortcomings of non-cryoprotective polysaccharides and retain considerable cell recovery rates for statistical significance testing. After medium exposure, the microplate was covered in parafilm and frozen at –80°C in a Styrofoam container designed to yield an optimal cooling rate of –1°C/min (±0.2°C). After 24 h, the microplate was thawed in a 37 °C water bath for 2 min. Each well content was diluted 3-fold with PBS and left to incubate at 37 °C, 5% CO_2_ for another 24 h, for cells to regain normal metabolic rate and to avoid an overestimation of viability due to post-thaw cellular apoptosis events that occur around the 6-hour mark post-thaw (Baust et al., 2009). Then, the liquid content was discarded, 100 µl of the metabolic indicator resazurin (0.08 mg/ml) were added and the microplate was left to incubate at 37 °C, 5% CO_2_ for 2 h. Absorbance was measured at 570 and 600 nm in a BioTek^®^ microplate reader and visualized in the BioTek^®^ Gen 5 software. Resazurin in the absence of cells was used as control of the experiment. The commercial cryogenic formulation CryoStor™ CS5 and 10% DMSO were used as positive control and DMEM was used as negative control. The post-thaw metabolic viability (PTV) data shown is the best possible fold-change obtained for each polysaccharide, regardless of concentration used, to reflect maximal performance for a given chemical structure. For histological data, a NIKON Eclipse Ti-S optical microscope connected to a NIKON D610 digital camera was used for image acquisition. The liquid contents of all wells were discarded and representative pictures were collected with the Camera Control Pro software.

### 2.4. Principal Component Analysis

To identify underlying patterns in the data that could explain the majority of the variation in post-thaw viability during cell cryopreservation using polysaccharides, principal component analysis (PCA) was performed. The dataset consisted of 26 discrete data points and 4 predictor variables: fucose (mol%), UA (mol%), neutral monomers (mol%) and molecular weight (MDa). Missing values were imputed as 0. A correlation matrix was generated to assess the presence of any high correlations among the predictor variables. A cumulative Scree plot of explained variance was generated to ascertain the optimal number of principal components (PC) to retain. A PC1– PC2 biplot with corresponding factor loadings expressed as variable eigenvectors was plotted to visualize the relationship between the variables and the principal components. Similar datapoints were clustered through K-means clustering, revealing an optimal 3-cluster system. A distance-to-the-mean plot was used to determine the optimal number of clusters. All analyses were computed using Python 3.9.5, using a combination Jupyter Notebook 6.1.4 and PyCharm Community Edition 2022.3.1 for code execution.

### 2.5. Statistical analysis

All data was collected as (at least) four replicate measurements, for each N=26 polysaccharides tested. Results are shown as mean ± σ^2^. Statistical significance was assessed with an ordinary ANOVA, with Sidak’s multiple comparisons test. Significance was reported using the NEJM *p*-value threshold nomenclature as follows, from ascending order of significance: 0.12 (ns), 0.033 (*), 0.002 (**), <0.001 (***), where ‘ns’ means not significant for a 95% confidence interval (CI), with α=0.05. Linear regression analyses are reported with *R^2^* and corresponding 95% CI. QSAR trend interpolations were represented by smoothing splines, a non-parametric analysis mediated by a goodness-of-fit penalty function that approximately fits quantitative data while avoiding overfitting. All statistical analysis was performed in GraphPad Prism 8.0.1.

## 3. Results

### 3.1. Metabolic post-thaw cell viability (PTV)

Adherent monolayer Vero cells were cryopreserved at –80 °C exposed to a freezing medium containing 10% DMSO and each test polysaccharide, at varying concentrations, dissolved in optimal DMEM culture medium. A list of the polysaccharides used and their compositional profile is shown in **Table 2**. **Figure 1** shows the average post-thaw viability (PTV) fold-change, relative to 10% DMSO, for every polysaccharide tested. The results were compared with the commercial cryoprotective formulation CryoStor™ CS5.

**Figure 1.**
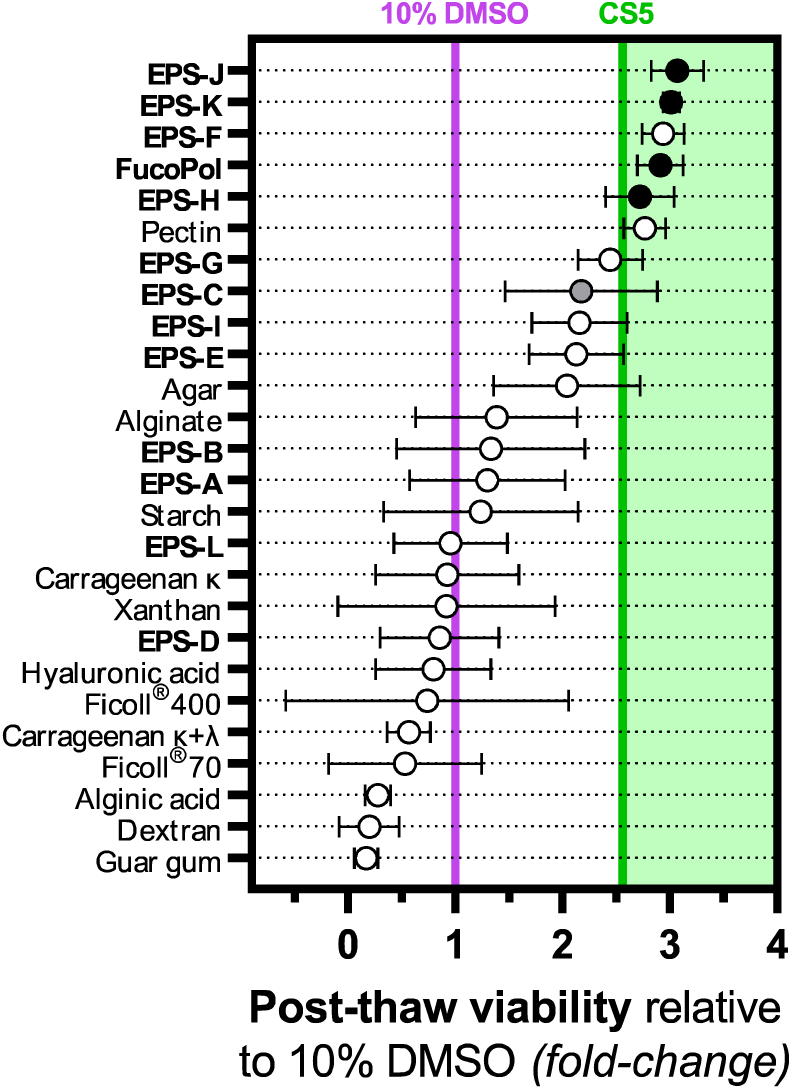
Ranked cryoprotective effect of the studied polysaccharides, relative to 10% DMSO performance (control). Single datapoints corresponding to each polysaccharide (N=4) are shown as white circles (○) with error bars, black circles (●) emphasize high-fucose content polysaccharides (above 20%), gray (●) gray indicates low fucose (1-5%) and white indicates no fucose. Vertical lines indicate threshold post-thaw metabolic viability for each control, purple for cells frozen in the sole presence of 10% DMSO and green for the commercial cryogenic formulation CryoStor™ CS5. Error bars are shown for replicate experiments (4≤N≤8).

**Table 2.** Polysaccharide dataset used in Vero cell cryopreservation experiments. The resulting biological post-thaw viability, compositional profile (mol%) and relevant ratios are shown (exact monomer compositions available in *SI*). PTV data is a replicate average (N=4–8) and is relative to 10% DMSO, which corresponds to a 1.0-fold change. Commercial polysaccharide data is limited to manufacturer information availability.

| EPS | PTV<br>(fold-change) | Fucose<br>(mol%) | Uronic Acids<br>(mol%) | MW<br>(MDa) | Fuc:UA<br>stoichiometry |
| --- | --- | --- | --- | --- | --- |
| ***Biotechnologically produced*** |  |  |  |  |  |
| **FucoPol** | 2.91 ± 0.90 | 34.0 ± 2.0 | 9.5 ± 0.5 | 3.75 ± 2.05 | 3.58 |
| **A** | 1.30 ± 0.14 | 0 | 42.1 | 3.1 ± 1.5 | 0 |
| **B** | 1.33 ± 0.50 | 0 | 45.5 | 4.6 | 0 |
| **C** | 2.17 ± 0.40 | 4.35 | 17.4 | 1.2 | 0.25 |
| **D** | 0.85 ± 0.12 | 0 | 45.0 | 1.4 | 0 |
| **E** | 2.13 ± 0.67 | 0 | 22.2 | 3.4 ± 0.9 | 0 |
| **F** | 2.94 ± 0.52 | 0 | 37.5 | 3.2 | 0 |
| **G** | 2.44 ± 0.63 | 0 | 50.0 | 0.51 ± 0.04 | 0 |
| **H** | 2.72 ± 0.41 | 37.0 | 23.0 | 4.2 | 1.61 |
| **I** | 2.15 ± 0.37 | 0 | n/d | 1.33 ± 0.57 | - |
| **J** | 3.07 ± 0.10 | 26.4 ± 0.03 | 9.3 ± 0.4 | 7.88 | 2.84 |
| **K** | 3.01 ± 0.67 | 45.0 ± 0.2 | 7.1 ± 0.8 | n/d | 6.36 |
| **L** | 0.95 ± 0.11 | 1.00 | 11.7 | n/d | 0.09 |
| ***Commercially sourced*** |  |  |  |  |  |
| **Pectin** | 2.77 ± 0.47 | 0 | 74 | 0.18 ± 0.13 |  |
| **Agar** | 2.04 ± 0.33 | 0 | 30 | 0.12 |  |
| **Alginate (G-rich)** | 1.38 ± 0.59 | 0 | 100 | 0.21 ± 0.19 |  |
| **Alginate (M-rich)** | 0.28 ± 0.09 | 0 | 100 | 0.22 |  |
| **Starch** | 1.24 ± 0.11 | 0 | 0 | 0.25 ± 0.24 |  |
| **Carrageenan κ** | 0.92 ± 0.25 | 0 | 0 | 0.25 |  |
| **Carrageenan κ+λ** | 0.57 ± 0.21 | 0 | 0 | 0.8 |  |
| **Xanthan** | 0.92 ± 0.32 | 0 | 20 | 2 |  |
| **Hyaluronic acid** | 0.79 ± 0.17 | 0 | 50 | 0.85 ± 0.35 |  |
| **Ficoll®400** | 0.74 ± 0.19 | 0 | 0 | 0.4 |  |
| **Ficoll®70** | 0.53 ± 0.11 | 0 | 0 | 0.07 |  |
| **Dextran** | 0.20 ± 0.08 | 0 | 0 | 0.04 ± 0.01 |  |
| **Guar** | 0.17 ± 0.07 | 0 | 0 | 0.22 |  |

\* 0 indicates a virtual ratio of 0, which compositionally groups UA-only polysaccharides into an average datapoint.

The freezing medium contained a mixture of each polysaccharide and 10% DMSO. Thus, PTV values near 1.0-fold indicate minimal to no cryoprotective effect for the given polysaccharide, and negative synergy below that. As the PTV of CryoStor™ CS5 is 2.56-fold, it was considered that for a polymer to be considered cryoprotective, it should at least differ by one standard deviation from the 10% DMSO benchmark (*i.e.,* a 2.0-fold change or above). By these standards, *agar*, *EPS-E*, *EPS-I*, *EPS-C* and *EPS-G* showed a mid-tier cryoprotective effect (PTV=2.04–2.44), while *pectin*, *EPS-H, FucoPol*, *EPS-F*, *EPS-K* and *EPS-J* showed high-tier cryoprotection that outperformed CryoStor™ CS5 (PTV=2.77–3.07).

#### 3.1.1. Influence of monomer composition

Interestingly, all polysaccharides with a high content in fucose (**Figure 1**, black circles) out-performed CryoStor™ CS5. *EPS-H* (37.0 mol% fucose) yielded a 2.7±0.3-fold change, *FucoPol* (34.0 ± 2.0 mol% fucose) yielded 2.9±0.2-fold, *EPS-K* (45.0 ± 0.2 mol% fucose) yielded 3.0±0.1-fold and *EPS-J* (26.4 ± 0.03 mol% fucose) yielded a 3.1±0.3-fold change (**Table 2**). In other words, all these polysaccharides outperformed CryoStor™ CS5 by 5.4, 13.2, 17.1 and 21.1%, respectively, in guaranteeing cell survival after thawing. Polysaccharides with a lower fucose content (**Figure 1**, gray circles) such as *EPS-*C (4.35 mol% fucose) and *EPS-L* (1.0 mol% fucose) show a relatively lower PTV, but appreciable enough to justify the observation of a relationship between fucose content and cryoprotective efficiency.

Polysaccharides with a high UA content still generated a high PTV, as is the case of *EPS-F* (PTV=2.94, 37.5 mol% UA) and *EPS-G* (PTV=2.44, 50 mol% UA) despite having no fucose in their composition (Figure SI.1), which agrees with previous observations of a relationship between polyanionicity and cryoprotective potential (Shtukenberg et al., 2017). Fucose and UA appear to be key features in ensuring high PTV, as their absence results in low-tier PTV, as is the case of *EPS-L*, which contains residual amounts of fucose (<1%), a comparatively low UA content of 11.7% (Table SI.1) and resulted in only a maximal PTV=0.95±0.11.

#### 3.1.2. Influence of structural factors

Slight exceptions in the relationship between monomer composition and PTV appear related to structural factors playing an additional role. For instance, *EPS-D* (no fucose, 45 mol% UA) resulted in a PTV=0.9±0.1 but has a similar composition to *EPS-G*, which had a PTV=2.4±0.6. The distinction is that *EPS-D* possesses a far lower zero-shear viscosity (*η*_0_=0.74 Pa·s) relative to all other polymers, which average at *ca.* 16 ≤ *η*_0_ ≤ 76 Pa·s (**Table SI.1**). It also contains a far lower MW of 1.4 MDa, while high-performing polysaccharides range between 3–7.8 MDa (**Table 2**).

*Pectin* (PTV=2.77±0.5) and *agar* (PTV=2.04±0.3) were the only commercial polysaccharides that showed an appreciable effect. Pectin has at least 74 mol% UA in its composition, 6.7 wt.% methoxy groups and is known to be hyperbranched (L. Chen et al., 2019). Agar is composed of linear agarose and an heterogenous mixture of agaropectins, which are also UA-rich, and is widely used for its gel-forming capabilities (Zeece, 2020). Two alginate polysaccharides of different G/M ratios were also tested, and despite their similarities, resulted in a significant difference in PTV outcome. Alginates are composed of α-(1→4) linked L-guluronic acid (GulA, or G) and D-mannuronic acid (ManA, or M) monomers (Skjåk-Bræk & Draget, 2012). The G-rich alginate (PTV=1.4±0.8, 75 mol% GulA) showed better performance than the M-rich alginic acid (PTV=0.28±0.12, 35 mol% GulA). Previous literature has shown that a higher GulA content generates steric hindrance around the –COOH group, introducing polymer folding and creating a more rigid structure (L. Chen et al., 2019; Yang et al., 2011). From a mechanical damage standpoint, polymer rigidity and high viscosity produce similar outcomes in what concerns beneficial osmoregulation (C. S. Wang et al., 2021; H. Wang et al., 2022) in a cryoprotection setting. Therefore, a cryoprotective outcome appears related to the overall ionic nature of monomer composition, whereas structural features like increased viscosity, conformation in solution and MW play a consequential complementary role.

### 3.2. Post-thaw cell histology

Cell histology is an equally important feature of cell healthiness alongside metabolic viability, and substances capable of preserving the native cell layer structure are of particular interest in real-world applications. **Figure 2** shows the microphotographs of Vero adherent cells after being cryopreserved in the presence of the five best performing polysaccharides, *EPS-J*, *EPS-K*, *EPS-F*, *FucoPol* and *EPS-H*, at each corresponding concentration that maximized PTV.

**Figure 2.**
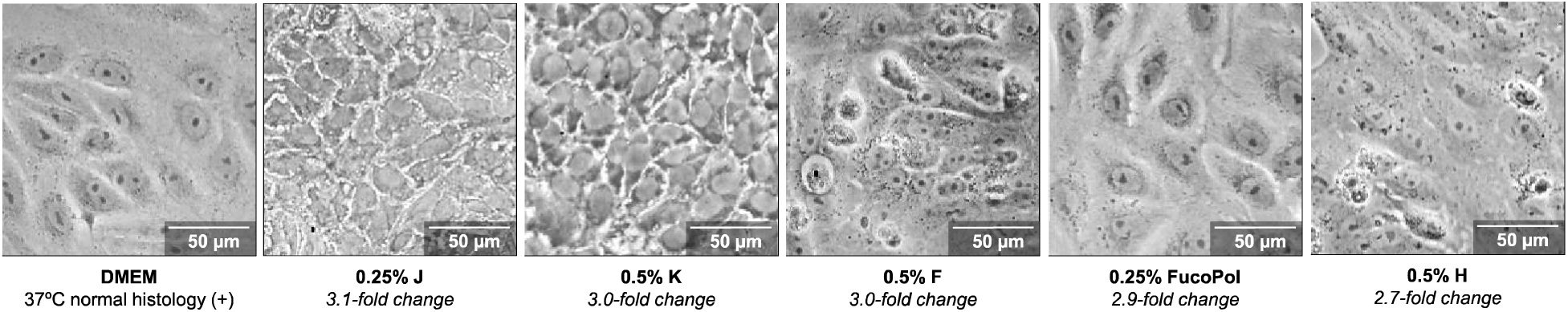
Histology of thawed Vero cells cryopreserved in polysaccharide freezing media. The top-five polysaccharides that generated the highest metabolic PTV in this study were selected and are labelled accordingly below each photomicrograph. A total magnification of 400× was used for image acquisition.

The morphology of Vero cells in DMEM culture medium is representative of normal histology and is characterized by adhered cells with cell-cell interactions conducing to a polygonal outstretched appearance. The nuclear and cytosolic contents are clearly distinguishable and cellular delimitations are perceivable from smooth outlines of each cell. Polysaccharide supplementation to the freezing medium resulted in diverse histological outcomes. The use of 0.25% *EPS-J* generates a more disorganized and compacted pattern of growth, with cells being slightly shrunk and showing high amounts of particulate matter (gray) near their outlines. The pattern is similar with 0.5% *EPS-K*, although a more discrete separation between each cell exists, cell size is similar to that of native Vero, but the nuclear contents are not visible. *EPS-H* and EPS-F resulted in the poorest adherence patterns, with some visible detachment, suggesting that during the 24-hour period post-thaw, cell structuration was most likely sub-optimal. Despite all polysaccharides resulting in high PTV, *FucoPol* was the only polysaccharide to demonstrate the ability to fully preserve native cell histology, with results nearly identical to that of Vero cells optimally grown at 37°C in DMEM. We suspect that the variance in cell morphology arises from each polysaccharide creating a characteristic polymeric matrix based on their individual structural features, which in turn results in novel patterns of cellular adherence without risking metabolic viability. In the case of FucoPol, slightly denser cell outlines can be observed with careful observation, suggesting that it reinforces the Vero natural adherence pattern but is still present in intercellular spacings.

In summary, *FucoPol* has demonstrated to be a model cryoprotective polysaccharide, optimizing both metabolic and histological viability, showing promising applicability in real-world formulations and distinguishing itself from other high-tier cryoprotective polysaccharides.

### 3.3. Qualitative split testing

The first step to deconvoluting the effects on cryoprotective outcome was to evaluate each variable considered to be of relevance separately. **Figure 3** shows comparative box plots that agglomerate PTV data based on fucose content, UA content and MW.

**Figure 3.**
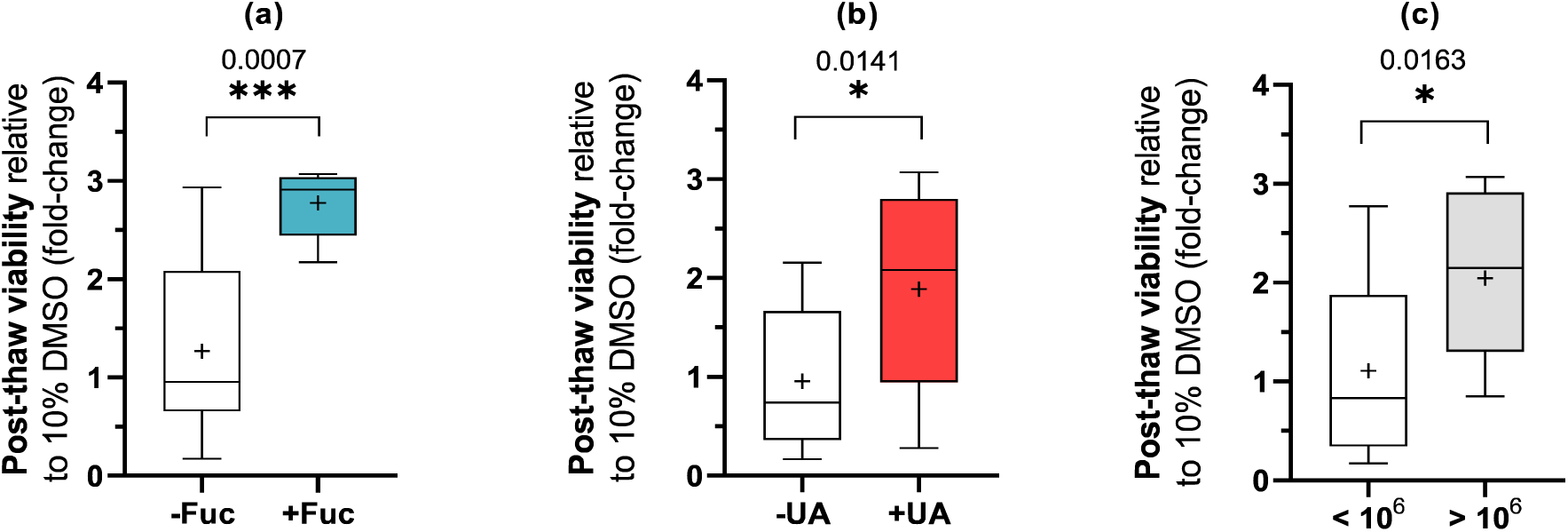
Single-variable effect distribution towards Vero cell post-thaw viability. Boxplots compare polysaccharides containing or not **(a)** fucose or **(b)** UA in their composition, and **(c)** having a molecular weight lower or higher than 1 MDa. Statistical significance between each comparison pair was calculated according to Sidak’s multiple comparisons test (* *p≤* 0.05, *** *p≤* 0.001).

Polysaccharides that contain any amount of fucose in their composition showed a mean PTV increase of 2.8±0.4-fold, compared to only 1.3±0.8-fold in its absence (**Figure 3a**, *p*=0.0007, N=26). This relationship strongly indicates that the presence of fucose is highly beneficial towards a positive cryoprotective outcome and results in high Vero cell viabilities after thawing. Polysaccharides containing any amount of UA in their composition showed a mean PTV increase of 1.9±0.9-fold (**Figure 3b**, *p*=0.0141, N=26), while in their absence the PTV dropped to that of the control (PTV=1.0±0.7), indicating the absence of cryoprotective function. The distribution of PTV data for UA-containing polysaccharides (red box) is broader than fucose-containing polysaccharides (cyan box) because fucose is mostly predominant in niche high-tier PTV polysaccharides, whilst UA presence is more ubiquitous in this dataset. Polysaccharides above 1 MDa showed a mean PTV increase of 2.1±0.8-fold. For smaller polysaccharides, the mean PTV was 1.1±0.9-fold (**Figure 3c**, *p*=0.0163, N=23). In general, cryoprotective polysaccharides possess a MW above 10^6^, and considering those for which PTV≥2.0, the observed high-performance average MW is 2.56±0.37 MDa, compared to an average 1.03±0.16 MDa for those non-cryoprotective. The difference between Ficoll^®^400 (PTV=0.74±1.32) and Ficoll^®^70 (PTV=0.53±0.71) of 400 and 70 kDa (**Table 1**), respectively, although nuanced, further accentuates this trend.

The dataset used in this study validates previously known trends in the literature, namely that high-molecular weight, negatively charged polymers have a greater cryoprotective potential than smaller, neutral polymers (Matsumura et al., 2022; Murray & Gibson, 2022). *Ipso facto*, the observed fucose content trend is equally valid. However, notice that high fucose-containing polysaccharides often had relatively low UA contents (average 12.2±6.3 mol%) compared to most UA-containing polysaccharides (average 39±28 mol%). So, alongside the corroboration of high MW (average 2.5×10^6^ Da) towards maximizing cryoprotective function, fucose appears to enhance the cryoprotective effect, but doing so at the expense of lower UA content. Instead of a predominant UA content that maximizes polyanionicity, the PTV data suggests that a stoichiometric balance exists, which maximizes function, and can be determined.

### 3.4. Quantitative structure-activity relationship (QSAR) analysis

Split testing has qualitatively shown that MW, fucose and UA contents are statistically contributive towards PTV outcome. **Figures 4–6** now quantitatively describe the effect of a given monomer on PTV outcome. Linear regression and spline interpolations applied to the discrete data can yield approximate spectral fingerprints that allow to infer optimal ranges for each variable that maximize cryoprotection.

**Figure 4.**
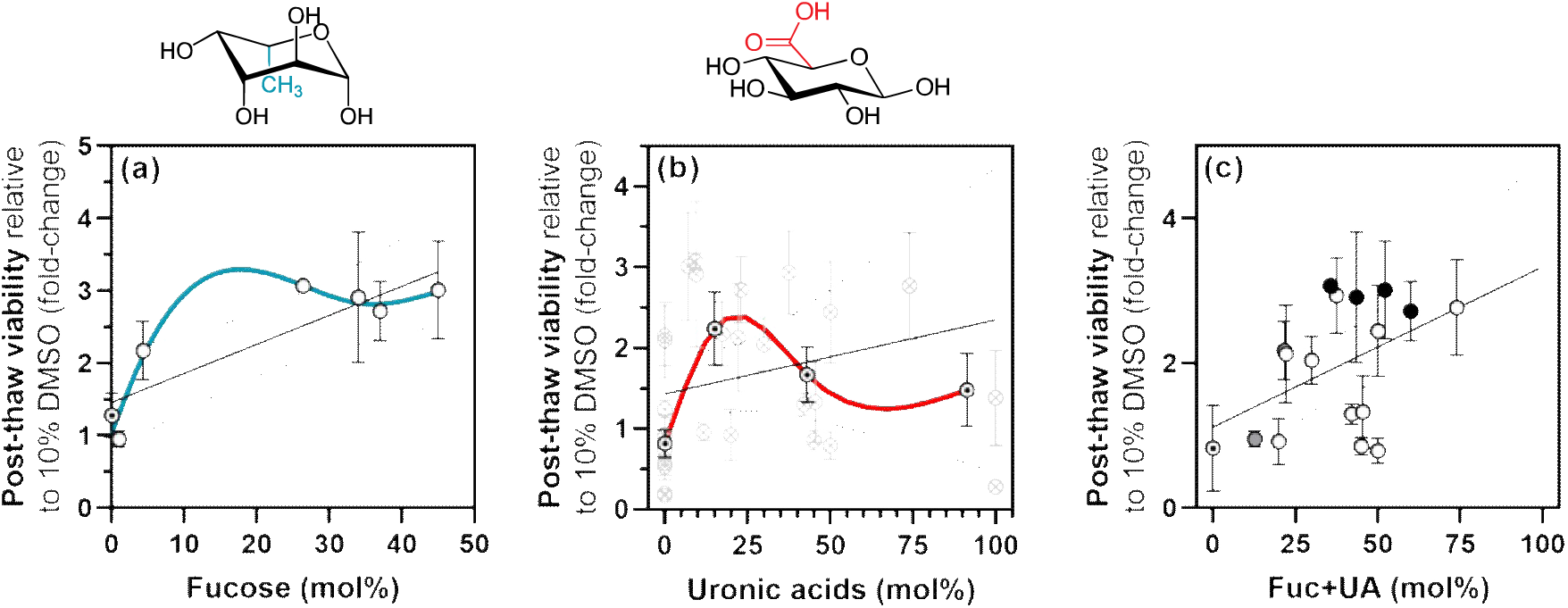
Influence of (a) fucose, (b) UA and (c) their combined contents on different polysaccharides in the PTV of cryopreserved Vero cells. Dotted circles (⨀) are composition-averaged datapoints. For visual legend, refer to the caption of Figure 1. For visual clarity, UA data was grouped into bins that correspond, from left to right, to concentration ranges of {0}, {0.1:25{, {25.1:50{ and {50-100} mol%. The relevant chemical structures above each plot, linear regression analysis and 95% CI are shown. Trend interpolation was performed with smoothing spline analysis and is shown as a bold cyan (fucose) or red (UA) line.

**Figure 5.**
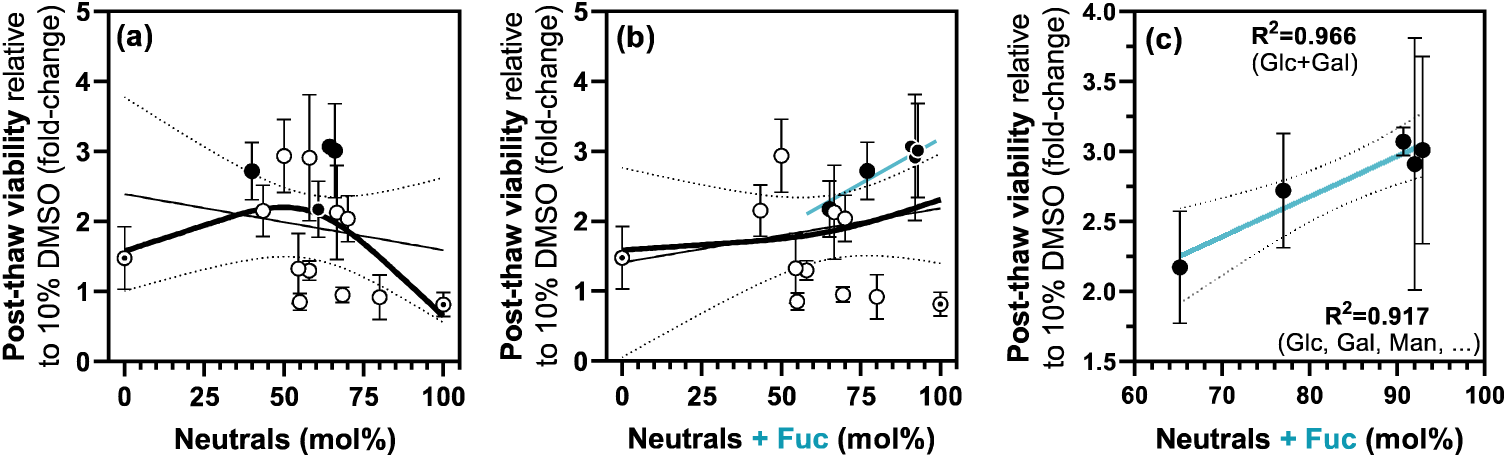
Influence of the combined neutral monomers content of different polysaccharides in the PTV of cryopreserved Vero cells. Dotted circles (⨀) are composition-averaged datapoints. For visual legend, refer to the caption of Figure 1. Scatter plots show the **(a)** neutral monomers content in glucose, galactose and mannose residues, **(b)** or also accounting for fucose as a neutral monomer. **(c)** Local linear regression of the contribution of fucose-containing polysaccharides towards PTV shown in (b). Linear regression analysis and 95% CI are shown, with corresponding *R*^2^ in inset. Trend interpolation was performed with smoothing spline analysis and is shown as bold black lines.

**Figure 6.**
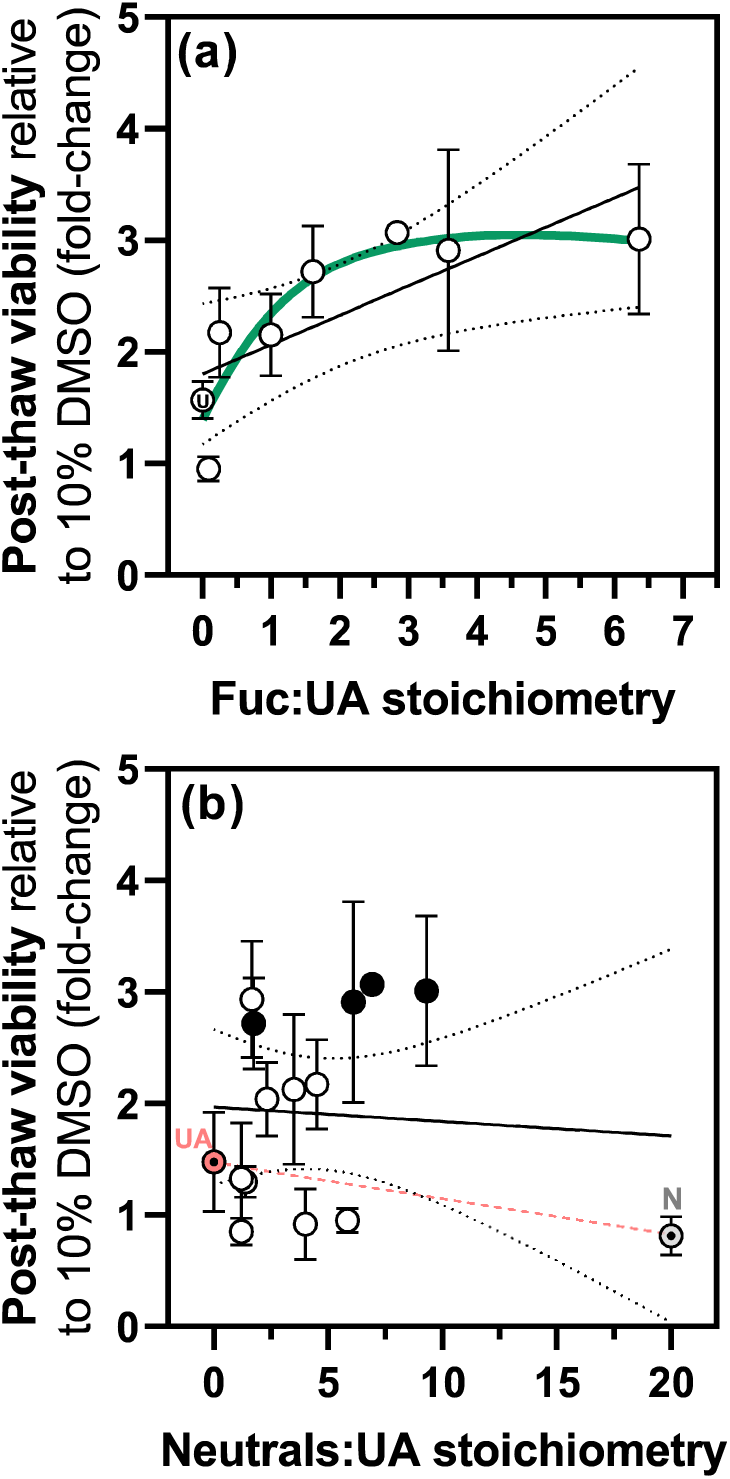
Influence of stoichiometric ratio between (a) fucose or (b) neutral monomers and the UA content on different polysaccharides in the post-thaw viability of cryopreserved Vero cells. Dotted circles (⨀) are composition-averaged datapoints. For visual legend, refer to the caption of Figure 1. In (a), ‘U’ corresponds to UA-only polysaccharides. In (b), the red ‘UA’ and gray ‘N’ datapoints correspond to UA-only and neutral polysaccharides, respectively, with the red dashed line emphasizing a PTV trend between both extremes. Linear regression analysis and 95% CI are shown. Trend interpolation was performed with smoothing spline analysis and is shown as a bold green or black line.

#### 3.4.1. Effect of fucose and uronic acids

Figure 4 shows the quantitative relationships between PTV and fucose content (Figure 4a), UA content (Figure 4b) and their combined fractions in the total compositional bulk (Figure 4c).

Fucose shows a positive association to PTV (Figure 4a). An increasing fucose fraction in cryoprotective polysaccharides boosts cryopreserved Vero cell viability, monotonically increasing at lower fucose contents and plateauing near PTV fold-changes of 2.9±0.1 and contents of 30–45 mol% (*R^2^*=0.768, N=7). In contrast, polysaccharides with no fucose resulted in an average PTV of 1.3±0.3, unbeknownst of their respective UA contents. This suggests that the presence of fucose in polysaccharides generates a strong enhancement of cryoprotective function by a factor of 2.3. A spline interpolation of the data suggests that the optimal fucose content peaks at *ca.* 18%, although the low peak height relative to fucose concentration 30 mol% and above suggest that a plateau is more likely (centered at *ca.* 35.7 mol%). At very high fucose concentrations (above 80 mol%), PTV enhancement is expected to decline due to a trade-off between fucose and other monomers. UA content in particular also has a beneficial effect on cryoprotective outcome (Figure 4b), as expected from negatively charged monomers (Matsumura et al., 2022), following a linear increasing trend (*R^2^*=0.202, N=24) until about 50 mol%, whereas an increase in UA stops generating a meaningful change in PTV. The absence of UA resulted in an average PTV fold-change of 1.0±0.2, indicating that polysaccharides not containing UA are not cryoprotective.

Conversely, their influence appears maximal between 9–26%, the latter value determined by spline interpolation. Therefore, low to mid-UA contents are preferred over high-UA contents (40+ mol%) towards maximizing PTV. The negligible contribution of very high UA contents towards PTV (PTV=1.7±0.3 for an average 43 mol% *vs*. PTV=1.5±0.5 for an average 91 mol%) also suggests that, like fucose, cryoprotective function is not maximized by maximizing negative formal charge. The combined contents of fucose and UA for the total composition of each polysaccharide (Figure 4c) also appears to have a decent predictive correlation towards cryoprotection (*R^2^*=0.212, N=26). A PTV increase is consistently observed in polysaccharides with higher absolute combined Fuc+UA fractions, hinting at their joint beneficial effect towards cryopreservation.

#### 3.4.2. Effect of neutral monomers

The interesting bioactive contribution of fucose towards PTV prompted comparison with monomers of equal chemical nature. Figure 5 contrasts the combined influence of ubiquitous glucose (Glc), galactose (Gal) and mannose (Man), as well as less frequent monomers (Rha, Fru; see **Table SI.1**), with that of fucose towards PTV outcome.

Figure 5a shows that exceeding amounts of either Glc, Gal or Man result in a prejudicial effect in PTV. The spline interpolation suggests that an optimal amount of neutral residues exists in mid-tier contents, peaking around 50 mol%, but the overall trend is of downwards concavity. However, when fucose is input into the neutral monomer content (Figure 5b), a trend inversion is observed. The cyan trendline shown in Figures 5b-c indicates that all high fucose-rich polysaccharides are clustered in a high PTV region, are mostly ordered by their increasing fucose content (from left to right: *EPS-C*, *EPS-H*, *EPS-J*, FucoPol, *EPS-K*) and establish a robust relationship with increasing PTV (*R^2^*=0.966, N=5). The strong contribution of fucose content towards reshaping the trendline of neutral monomer influence in PTV not only reinforces previous observations that fucose plays a significant role in enhancing cryoprotective outcome, but also that its chemical nature must differ from all other neutral monosaccharides.

#### 3.4.3. Optimal stoichiometric ratios

Using an absolute QSAR analysis, the fractional ranges of fucose, UA and neutral monomers which optimize PTV were obtained. A combinatorial average of optimal fucose (18– 35.7 mol%), optimal UA (9–26 mol%) and complementary neutral monomer (40–48 mol%) ranges yielded an average 98.5±6.6% total osidic fraction, which roughly approximates to the complete chain fraction described by chemical composition (100.0±5.5%). These ranges prove to be good absolute estimates for optimal PTV, but we can further define relativistic optimal stoichiometric ratios. Figure 6 shows the effect of increasing neutral monomer or fucose contents, with respect to a normalized UA content.

Figure 6a shows that a gradual increase in the fucose content of a given polysaccharide relative to its UA content generates a consistent increase in PTV outcome after cryopreservation (*R^2^*=0.585, N=14). This observation agrees with a PTV-enhancing effect of fucose. The highest ratio datapoint (Fuc:UA*=*6.4) corresponds to *EPS-K*, a polysaccharide with PTV=3.0±0.7. A quasi-linear correlation is observed for Fuc:UA ratios between 0–2 and a plateau at PTV=3 is achieved for higher ratios. The presence of fucose, either in sub-or supra-UA amounts, generates a perceivable PTV increase, just from the fact that increasing fucose is present, rather than other neutral monomers. When Fuc:UA ≥ 2, the linearity with PTV breaks as greater absolute amounts of fucose will not outweigh the beneficial effects of negative formal charge. Conversely, Figure 6b shows that an increase in neutral monomer content (mostly Glc, Gal, Man) relative to constant UA content is mostly decoupled from a change in PTV (*R^2^*=0.005, N=20). At compositional boundaries, UA-only polysaccharides resulted in PTV=1.9±0.5, compared to those fully neutral (PTV=1.1±0.3).

In summary, stoichiometric trends validate the observations that (i) fucose has a chemically different contribution compared to other neutral monomers and (ii) it enhances the cryoprotective effect of UA-based polysaccharides. The four polysaccharides of highest Fuc:UA stoichiometry (6.4 for *EPS-K*, 3.6 for *FucoPo*l, 2.8 for *EPS-J*, 1.6 for *EPS-H*) are precisely those of high-fucose (45, 34, 26.4, 37 mol%), mid-UA composition (7.1, 9.5, 9.3, 23 mol%), respectively. The decoupled effect of neutral monomers from PTV suggests their presence in a polysaccharide to be solely of linker character. However, no single constituent appears expendable: PTV is optimized with a Fuc:UA stoichiometric balance, rather than with fucoidan-or uronan-type polysaccharides.

### 3.5. Principal Component Analysis

The QSAR compositional spectrum for optimal cryoprotective performance during Vero cell cryopreservation for the polysaccharide dataset studied is summarized in Figure 7a. The UA (9–26 mol%) and fucose (18–35.7 mol%) optimal ranges previously discussed are present, alongside the slightly negative change in PTV with increasing neutral content. Polysaccharides that do not contain fucose and UA (*y-*intercept) demonstrate a PTV indistinguishable from the 10% DMSO control group (1.0-fold) supplemented in all experiments, indicating the absence of function. A slight increase in UA composition to 5–10 mol% generates a 60–100% PTV increase, corroborating the strong influence of polyanionicity in cryopreservation settings, easily outweighing the cryoprotective effect of 10% DMSO by a significant 2-fold. However, it is only in the combined presence of UA and fucose that a polysaccharide can competitively outperform the cryoprotective effect of CryoStor™ CS5 (2.6-fold). Such performance is observed in a broad fucose content range (8–30 mol%), suggesting the robustness of fucose-rich polysaccharides of variable composition in ensuring greater cell survival. Above 30 mol% UA, the potential detriment in PTV outcome is levelled by an increased fucose contribution (*see box emphasis*).

**Figure 7.**
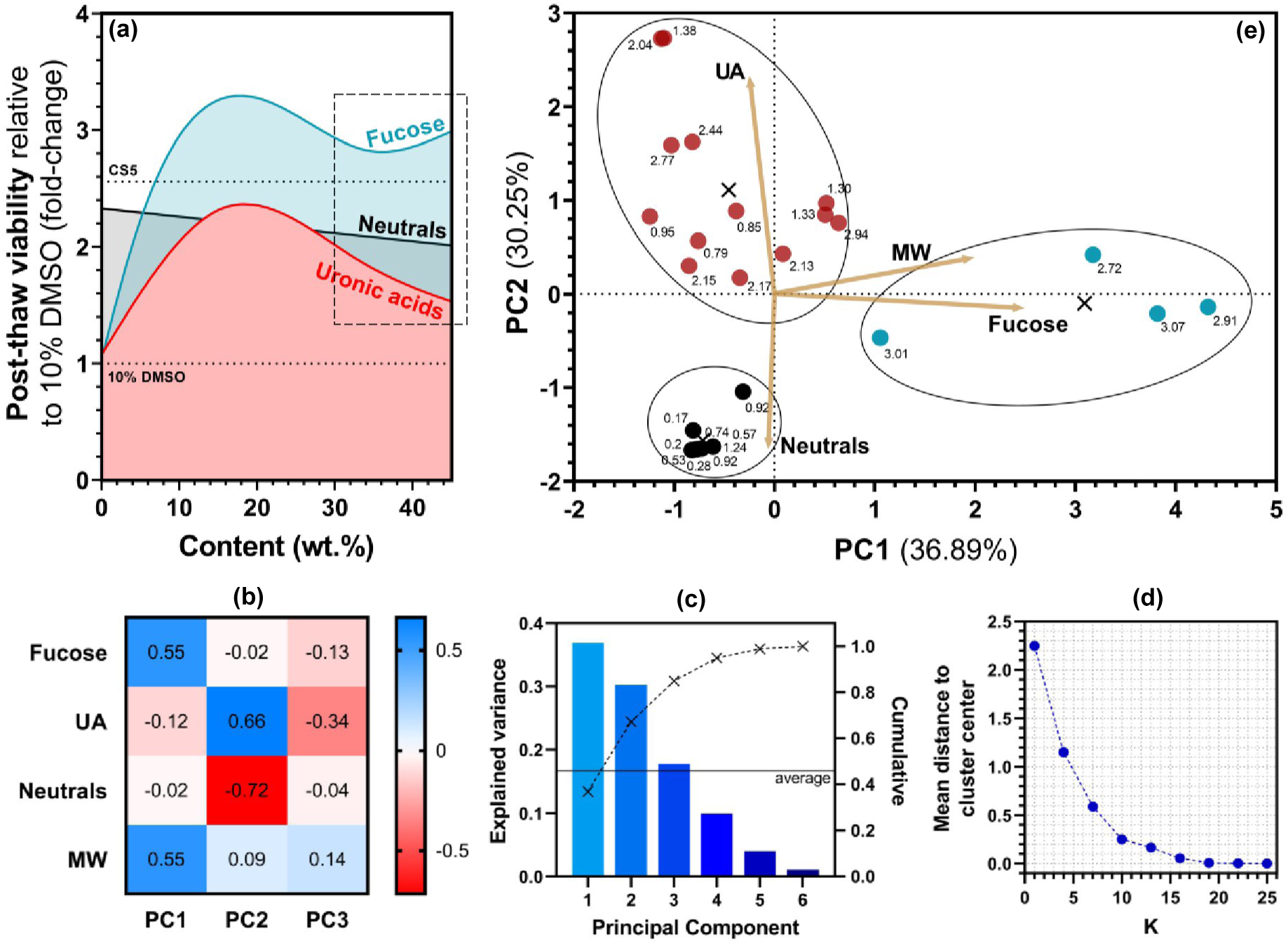
Summary of multivariable effect deconvolution in cryoprotective polysaccharides. (a) QSAR optimal compositional spectrum showing the optimal range for fucose and UA content, and comparative groups: control (10% DMSO) and commercial cryogenic formulation (CryoStor™ CS5); **(b)** PCA biplot with eigenvector loadings for each predictor variable and point clustering. The arrows that extend outward from the origin are eigenvectors, which indicate the variance loading that a variable has on each principal component. The degree of influence that each element has on a principal component is visualized by the angle and length of the arrow relative to the axes. **(c)** Correlation matrix, quantitatively describing the contribution of each variable to a principal component. **(d)** Scree plot of explained variance (bars), complemented by the cumulative explained variance for all principal components (dashed line). **(e)** *K*-means clustering score model that iteratively calculates the mean distance of each datapoint to *N* cluster centers and returns an optimal *K* clusters.

The maximization of cryoprotective function in polysaccharides then appears based on a compositional compromise between negatively charged species (UA) known for their polar interaction with water and disruptive effect on ice physics, and the neutral monomer fucose. The importance of the latter given its nature hinted at a mechanism of action different from the presupposed polyanionicity. Thus, we performed a principal component analysis (PCA), to assess how each predictor variable independently explained a change in PTV outcome (Figures 7b**–e**). By decomposing the individual contributions of each variable, similar performing predictors were highlighted by *k*-means clustering, and each principal component (PC) was attributed to a plausible mechanism of action involved in the overall biological expression of cryopreservation.

First, the correlation matrix in Figure 7b and the Scree plot in Figure 7c show that most of the cumulative variance in PTV can be explained by two major PCs, where PC1 has major contributions from fucose content (+0.55) and molecular weight (+0.55), whilst PC2 is mostly constituted by opposing contributions between UA (+0.66) and neutral monomers (–0.72). A third PC (17.8%) did not significantly differ from the mean explained variance (Figure 7c, 16.7%) and was disregarded. Second, a *k*-means clustering algorithm computed the distance of each PTV datapoint to a centroid average (Figure 7d) and determined that PTV datapoints can be split in three different clusters of data. Lastly, the biplot shown in Figure 7e factors in the correlation data as variable eigenvectors that constitute each PC, thus representing how a combination of variables may contribute to a given mechanism of action.

In summary, the variability in cryopreservation outcomes arise as a combination of two mechanisms: one mediated by opposing chemical charge of anionic UA *vs*. neutral monomers (PC2); and another pointing to a MW-scaled mechanism exclusively mediated by fucose (PC1). Although a higher MW synergizes with increased UA content (+0.09), as is the case of high MW (3.2 MDa), high PTV (2.94±0.52) fucose-free *EPS-F,* the effect is much more noticeable with fucose content (+0.55). Notice the negligible contribution of fucose and UA outside their corresponding PCs (Figure 7b), suggesting these monomers act by different mechanisms. The resulting three clusters originate from high PTV outcomes obtained due to high polyanionicity (red) or high-fucose regimes (cyan), and low PTV outcomes associated to high neutrality (black).

## 4. Discussion

It is widely accepted that cryoprotective action is related to the presence of formal charge. Molecular systems with the ability of charge delocalization or transfer can interact with water molecules and disrupt ice growth (Shtukenberg et al., 2017). In polymers, this translates to polyanionicity and benefits from a cumulative increase in molecular weight (Shtukenberg et al., 2017). Cold-adapted microorganisms often produce EPS with a predominantly high UA content relative to other extremophiles (J. Wang et al., 2019). However, the established assumption is that the contribution of increased negative charge towards molecular interactions with unfrozen water follows a linear trend. This presumes uronan polysaccharides will maximize cryoprotective function. Here, we have shown that this is not true (*e.g.* alginates, pectin), as UA content is optimal in the 9–26 mol% range, followed by a decrease in PTV outcome and a plateau from 40 mol% onwards. This observation aligns with some known mechanisms that might explain the non-linear behavior. Initially, a linearly increasing PTV outcome with UA content correlates to a contribution of increasing negative charges in the polymeric chain establishing more polymer-water and polymer-ice interactions (Ewart et al., 1999; Kratochvílová et al., 2016). Above a certain UA threshold, polymer-polymer chain interactions start dominating the system and may result in polymer aggregation (Engel et al., 2004; Trachtenberg, 1986), chain entanglement, cross-linking (Ganesan et al., 2016; Torres et al., 2015) and enhanced water structuration (Grossutti & Dutcher, 2016). Naturally, the magnitude or emergence of these effects will vary based on the overall composition and structure. In this scenario, any cryoprotective effect will be dominated by diffusion-hindered processes rather than chemical binding that leads to crystalline disruption, and thus will not scale with chemical composition.

Due to the influence of formal charge in function, the presence of neutral monosaccharides in the structure has often been overlooked and sidelined as a default linker role in the polymer chain. The observation that glucose, galactose and mannose are ubiquitous in polysaccharides from diverse adaptive habitats (J. Wang et al., 2019) has further solidified this belief. However, in this work, we argue that fucose has a chemical contribution distinguishable from its other neutral monomer counterparts despite its chemical neutrality, disputing the governing assumption that polyanionicity is the sole dominant effect needed to explain cryoprotection. We observed that not only does an increasing fucose content maximize PTV, optimally ranging between 18–35.7 mol%, but it also follows a similar trend to UA. There appears to exist a beneficial trade-off between having an appropriate charge balance of UA interleaved by neutral fucose, rather than only UA in the composition. In truth, the presence of fucose is prevalent in polysaccharides from cold-adapted species and regularly co-exists with some amount of UA (J. Wang et al., 2019). The systematic observation of an interplay between fucose and UA manifesting a key role in polysaccharide cryoprotection is, to the best of our knowledge, a novel finding, and has two main implications. First, it demonstrates that cryoprotective outcome is maximized neither by fucose nor UA alone but relies on a chemical balance between both residues. This observation resembles what was reported for the ice recrystallization inhibition (IRI) mechanisms of polyampholytes (Stubbs et al., 2017), where maximal IRI activity is not obtained by maximizing polar moieties but by designing polymers with alternating hydrophilic and hydrophilic blocks, because this tacticity mimics the intermolecular spacings of a growing ice facet, boosting interfacial affinity. Second, the neutral nature of fucose might imply the existence of a mechanism that differs from charge contribution, a hypothesis supported by PCA and whose physical nature we expand upon hereafter.

Two principal components explained the variability in PTV. One was governed by the presence or absence of negative charges (UA *vs.* neutral monomers), hinting at an electrostatic mechanism related to ice physics disruption. The other reflected a mechanism strongly dominated by fucose presence that scaled with MW, which we argue might be associated with cell membrane stabilization effects. In fact, there are extensive accounts in the literature that point to fucose having a strong biological correlation to cell membrane interactions that are beneficial in cryobiology. Fucose plays a major role in cell recognition (Schneider et al., 2017), membrane binding (Ma et al., 2006), cellular homeostasis (Listinsky et al., 1998) and inflammatory processes (Y. Li et al., 2021), regulating the binding of carbohydrate ligands to selectins (Listinsky et al., 1998), fucose-mannose receptors (Péterszegi, Fodil-Bourahla, et al., 2003), fucose-recognizing lectins (Péterszegi, Fodil-Bourahla, et al., 2003) and specific CD16 surface receptors (e.g. FcγRIIIa) (Schneider et al., 2017). It is suspected that its hydrophilic-hydrophobic nature is responsible for its function as a specific recognition site in some vertebrate hemagglutinins (Thomas et al., 1956). The cell surface is often histochemically marked with α-L-fucose to checkpoint specific events, such as the maturation of lectins in human epidermal cells (Hietanen & Salo, 1984) which progresses whenever the cell surface contains α-L-fucose (Zieske & Bernstein, 1982). Moreover, it participates in restoring the levels of epithelial tight junction proteins (Y. Li et al., 2021), calcium chelation (Listinsky et al., 1998), proliferation of fibroblasts (Péterszegi, Isnard, et al., 2003), wound healing (Ye & Azar, 1998) and antioxidant ROS scavenging (Péterszegi, Robert, et al., 2003). The recovery of a cell from freezing stressors after thawing depends on all these factors, which are rooted on (and contribute to) homeostatic regain. In polysaccharides, fucose has early phylogenetic appearances in algae and fungi (Péterszegi, Fodil-Bourahla, et al., 2003), being sulphated in some instances as in fucoidans (B. Li et al., 2008) but, to the best of our knowledge, its in-chain importance in cryoprotection has not been elucidated before. Some functional parallelisms can be drawn from the multifunctionality previously described for the fucose-containing polysaccharide FucoPol, such as accounts of the salt tolerance properties of fucoidan (Zou et al. 2020) and similar antioxidant effect of fucose-rich polysaccharides (Péterszegi et 2002), but the literature on fucose-rich cryoprotectants is scarce.

We observed that the high-performing cryoprotective polysaccharides showed extracellular matrix-like scaffolding properties, promoting the growth of metabolically viable Vero cells in unique patterns of adherence. EPS-producing microorganisms usually inhabit an EPS scaffold that facilitates nutrient diffusion, offers protection from external stressors and promotes cell-cell communication (Nwodo et al., 2012). In particular, psychrophilic EPS that are responsible for protecting cells against freezing-related injury by disrupting ice growth and attenuating osmotic stress (Deming & Young, 2017), contain oddly high amounts of fucose (J. Wang et al., 2019). So, their perceivably strong contribution towards Vero cell adherence may reflect a leveraging of the membrane interaction capacity of EPS, finetuned by design in their composition and structure, thus explaining the prominence and benefits of fucose in a cryopreservation setting.

The ability of fucose to strongly mediate cell membrane interactions might arise from several factors. First, its L-configuration enables distinct stereochemical properties to arise (Flowers, 1981). Contrary to its parent monomer D-galactose, the absence of an –OH group at C-6 enables neighboring residues to be more axially distributed, thus sterically more available (Péterszegi, Fodil-Bourahla, et al., 2003), and the methyl group at C-5 (Figure 4a) introduces a potentially beneficial amphipathic character, as mentioned before (Stubbs et al., 2017). Second, although it only appears in terminal positions of vertebrate glycoconjugates (Russell et al., 1998), it emerges regularly along the chain of bacterial and algal polysaccharides (J. Wang et al., 2019), increasing the number of available *loci* for membrane interaction. This might explain the co-influence of MW and fucose in PC1: the fact they integrate as predictors of the same mechanism is most likely because the number of interaction points with the cell membrane scales with a bigger polymer chain. Therefore, the plausible introduction of chain folding by L-type isomers (Flowers, 1981), at the expense of a lesser amount of UA, but increased axial distribution of their neighboring –COOH (Figure 4b) (Péterszegi, Fodil-Bourahla, et al., 2003) and increased hydrophobic character (Stubbs et al., 2017) that potentiates cell membrane interactions (Ma et al., 2006), could explain the enhanced cryoprotective capabilities of polymers rich in L-fucose. L-rhamnose (Rha) has also shown membrane interaction-based cryoprotection, albeit much lower than fucose (Santarius & Bauer, 1983) but its peculiar stereochemistry may enable increased axial negative charge exposure, which might explain the high PTV values observed for the fucose-free *EPS-E* (PTV=2.1±0.7, 11.1 mol% Rha) and *EPS-F* (PTV=2.9±0.5, 12.5 mol% Rha).

Ultimately, the presence of fucose in a high-MW polysaccharide plays a critical role in enhancing cellular cryopreservation outcomes. However, cryoprotection is usually an umbrella term that nests phenomena like polymer-ice interactions and membrane stabilization effects that contribute to cell survival in a convoluted way, and distinction between both is often deemed unnecessary. By dissecting cryopreservation causality, we have shown that a successful outcome arises from a more complex set of predictors than simply polyanionicity. We hypothesize that a sequential pattern of fucose interleaved by UAs, much like the structural repeating unit of FucoPol (Guerreiro et al., 2020), maximizes PTV because it provides the optimal stoichiometric equilibrium towards enhancing molecular interactions on two fronts: (i) with the water molecules at the *quasi*-liquid layer of a growing ice crystal front and (ii) with the cell membrane. Understanding this causal relationship is important for the intentional design of promising polysaccharide-based cryoprotective formulations with enhanced potential, which can provide a safer, greener, biocompatible alternative to the use of glycerol and DMSO as cryoprotectants.

## 5. Conclusion

This work has demonstrated the critical role that in-chain fucose plays in the viability enhancement of Vero cells cryopreserved in polysaccharide media containing 10% DMSO. Vero cells experienced average metabolic PTV fold-changes of 0.8-fold for non-cryoprotective, 2.2-fold for cryoprotective and 2.8-fold for high fucose-containing polysaccharides, up to 3.1-fold, showing that some polysaccharides can outperform the optimized commercial cryopreservation formula CryoStor™ CS5 (2.6-fold). Cryoprotective performance is maximal when polysaccharides above 1 MDa have a high-fucose (18–35.7 mol%), mid-UA (13.5–26 mol%) compositional fingerprint. Optimal PTV was achieved with a Fuc:UA stoichiometric balance of 1.5:1 and higher, rather than with the dominance of one feature, challenging the assumption of polyanionicity as sole predictor of cryoprotection. Two major mechanisms explained the variability in PTV: charge-dominated ice growth disruption (UA *vs.* neutrals) and membrane stabilization effects (fucose, MW-scaled). Fucose-containing, UA-containing polysaccharides may actively contribute to polymer-water and polymer-cell interactions, disrupting ice growth in the former and minimizing lethal volumetric fluctuations in the latter. The study of structure-function relationships has enabled the mapping of optimal outcomes to predictor variables. By leveraging the high-throughput and flexibility of the bacterial machinery in biotechnology, the development of low-cost, bio-based, high-performance cryoprotective formulations can be accelerated by intentional design. Further studies are under way to confirm the role of polymeric fucose in cell membrane stabilization during freezing, and quantify membrane integrity alongside metabolic viability, in attempts to validate the existence of a membrane-associated mechanism of action.

## Author contributions

B.M.G. conceptualized study, performed experiments, data analysis, wrote manuscript. P.C-R. provided resources, wrote manuscript, reviewed manuscript. F.M. and H.P. performed experiments, wrote manuscript. X.M and J.G. provided resources and reviewed manuscript; F.F. conceptualized study, reviewed manuscript, supervised. J.L. and J.S. reviewed manuscript, supervised. All authors have read and agreed to the published version of the manuscript.

## Data Availability Statement

The data presented in this study is available on request from first authors or corresponding authors.

## References

1. Akan, G., & Oner, E. T. (2021). Gel Properties of Microbial Polysaccharides. Polysaccharides of Microbial Origin, 1–20.

2. Araújo, D., Martins, M., Concórdio-Reis, P., Roma-Rodrigues, C., Morais, M., Alves, V. D., Fernandes, A. R., & Freitas, F. (2023). Novel Hydrogel Membranes Based on the Bacterial Polysaccharide FucoPol: Design, Characterization and Biological Properties. Pharmaceuticals 2023, Vol. 16, Page 991, 16(7), 991.

3. Baptista, S., Pereira, J. R., Gil, C. V., Torres, C. A. V., Reis, M. A. M., & Freitas, F. (2022). Development of Olive Oil and α-Tocopherol Containing Emulsions Stabilized by FucoPol: Rheological and Textural Analyses. Polymers 2022, Vol. 14, Page 2349, 14(12), 2349.

4. Baust, J. G., Gao, D., & Baust, J. M. (2009). Cryopreservation: An emerging paradigm change. Organogenesis, 5(3), 90–96.

5. Benson, J. D., Higgins, A. Z., Desai, K., & Eroglu, A. (2018). A toxicity cost function approach to optimal CPA equilibration in tissues. Cryobiology, 80, 144.

6. Bojic, S., Murray, A., Bentley, B. L., Spindler, R., Pawlik, P., Cordeiro, J. L., Bauer, R., & de Magalhães, J. P. (2021). Winter is coming: the future of cryopreservation. BMC Biology 2021 19:1, 19(1), 1–20.

7. Carillo, S., Casillo, A., Pieretti, G., Parrilli, E., Sannino, F., Bayer-Giraldi, M., Cosconati, S., Novellino, E., Ewert, M., Deming, J. W., Lanzetta, R., Marino, G., Parrilli, M., Randazzo, A., Tutino, M. L., & Corsaro, M. M. (2015). A unique capsular polysaccharide structure from the psychrophilic marine bacterium Colwellia psychrerythraea 34H that mimics antifreeze (glyco)proteins. Journal of the American Chemical Society, 137(1), 179–189.

8. Carillo, S., Pieretti, G., Parrilli, E., Tutino, M. L., Gemma, S., Molteni, M., Lanzetta, R., Parrilli, M., & Corsaro, M. M. (2011). Structural investigation and biological activity of the lipooligosaccharide from the psychrophilic bacterium Pseudoalteromonas haloplanktis TAB 23. Chemistry (Weinheim an Der Bergstrasse, Germany), 17(25), 7053–7060.

9. Casillo, A., Fabozzi, A., Russo Krauss, I., Parrilli, E., Biggs, C. I., Gibson, M. I., Lanzetta, R., Appavou, M. S., Radulescu, A., Tutino, M. L., Paduano, L., & Corsaro, M. M. (2021). Physicochemical Approach to Understanding the Structure, Conformation, and Activity of Mannan Polysaccharides. Biomacromolecules, 22(4), 1445–1457.

10. Casillo, A., Parrilli, E., Sannino, F., Mitchell, D. E., Gibson, M. I., Marino, G., Lanzetta, R., Parrilli, M., Cosconati, S., Novellino, E., Randazzo, A., Tutino, M. L., & Corsaro, M. M. (2017). Structure-activity relationship of the exopolysaccharide from a psychrophilic bacterium: A strategy for cryoprotection. Carbohydrate Polymers, 156, 364–371.

11. Chen, L., Ge, M. D., Zhu, Y. J., Song, Y., Cheung, P. C. K., Zhang, B. B., & Liu, L. M. (2019). Structure, bioactivity and applications of natural hyperbranched polysaccharides. Carbohydrate Polymers, 223.

12. Chen, X., Zhang, Y., Han, Y., Li, Q., Wu, L., Zhang, J., Zhong, X., Xie, J., Shao, S., Zhang, Y., & Wu, Z. (2019). Emulsifying Properties of Polysaccharide Conjugates Prepared from Chin-Brick Tea. Journal of Agricultural and Food Chemistry, 67(36), 10165–10173.

13. Concórdio-Reis, P., Alves, V. D., Moppert, X., Guézennec, J., Freitas, F., & Reis, M. A. M. (2021). Characterization and Biotechnological Potential of Extracellular Polysaccharides Synthesized by Alteromonas Strains Isolated from French Polynesia Marine Environments. Marine Drugs 2021, Vol. 19, Page 522, 19(9), 522.

14. Concórdio-Reis, P., David, H., Reis, M. A. M., Amorim, A., & Freitas, F. (2023). Bioprospecting for new exopolysaccharide-producing microalgae of marine origin. International Microbiology, 1–8.

15. Concórdio-Reis, P., Pereira, J. R., Torres, C. A. V., Sevrin, C., Grandfils, C., & Freitas, F. (2018). Effect of mono– and dipotassium phosphate concentration on extracellular polysaccharide production by the bacterium Enterobacter A47. Process Biochemistry, 75, 16–21.

16. Concórdio-Reis, P., Pereira, C. V., Batista, M. P., Sevrin, C., Grandfils, C., Marques, A. C., Fortunato, E., Gaspar, F. B., Matias, A. A., Freitas, F., & Reis, M. A. M. (2020). Silver nanocomposites based on the bacterial fucose-rich polysaccharide secreted by Enterobacter A47 for wound dressing applications: Synthesis, characterization and in vitro bioactivity. International Journal of Biological Macromolecules, 163, 959–969.

17. Concórdio-Reis, P., Serafim, B., Pereira, J. R., Moppert, X., Guézennec, J., Reis, M. A. M., & Freitas, F. (2023). Exopolysaccharide production by the marine bacterium Alteromonas macleodii Mo169 using fruit pulp waste as the sole carbon source. Environmental Technology & Innovation, 30, 103090.

18. Cui, R., Zhang, L., Ou, R., Xu, Y., Xu, L., Zhan, X. Y., & Li, D. (2022). Polysaccharide-Based Hydrogels for Wound Dressing: Design Considerations and Clinical Applications. Frontiers in Bioengineering and Biotechnology, 10.

19. Deming, J. W., & Young, J. N. (2017). The role of exopolysaccharides in microbial adaptation to cold habitats. Psychrophiles: From Biodiversity to Biotechnology: Second Edition, 259–284.

20. Engel, A., Thoms, S., Riabesell, U., Rochelle-Newall, E., & Zondervan, I. (2004). Polysaccharide aggregation as a potential sink of marine dissolved organic carbon. Nature 2004 428:6986, 428(6986), 929–932.

21. Ewart, K. V., Lin, Q., & Hew, C. L. (1999). Structure, function and evolution of antifreeze proteins. Cellular and Molecular Life Sciences: CMLS, 55(2), 271–283.

22. Fialho, L., Araújo, D., Alves, V. D., Roma-Rodrigues, C., Baptista, P. V., Fernandes, A. R., Freitas, F., & Reis, M. A. M. (2019). Cation-mediated gelation of the fucose-rich polysaccharide FucoPol: preparation and characterization of hydrogel beads and their cytotoxicity assessment. International Journal of Polymeric Materials and Polymeric Biomaterials, 1–10.

23. Flowers, H. M. (1981). Chemistry and biochemistry of D– and L-fucose. Advances in Carbohydrate Chemistry and Biochemistry, 39(C), 279–345.

24. Freitas, F., Alves, V. D., & Reis, M. A. M. (2011). Advances in bacterial exopolysaccharides: from production to biotechnological applications. Trends in Biotechnology, 29(8), 388–398.

25. Ganesan, M., Knier, S., Younger, J. G., & Solomon, M. J. (2016). Associative and Entanglement Contributions to the Solution Rheology of a Bacterial Polysaccharide. Macromolecules, 49(21), 8313–8321.

26. Gilfanova, R., Callegari, A., Childs, A., Yang, G., Luarca, M., Gutierrez, A. G., Medina, K. I., Mai, J., Hui, A., Kline, M., Wei, X., Norris, P. J., & Muench, M. O. (2021). A bioinspired and chemically defined alternative to dimethyl sulfoxide for the cryopreservation of human hematopoietic stem cells. Bone Marrow Transplantation 2021 56:11, 56(11), 2644–2650.

27. Giwa, S., Lewis, J. K., Alvarez, L., Langer, R., Roth, A. E., Church, G. M., Markmann, J. F., Sachs, D. H., Chandraker, A., Wertheim, J. A., Rothblatt, M., Boyden, E. S., Eidbo, E., Lee, W. P. A., Pomahac, B., Brandacher, G., Weinstock, D. M., Elliott, G., Nelson, D., … Toner, M. (2017). The promise of organ and tissue preservation to transform medicine. Nature Biotechnology 2017 35:6, 35(6), 530–542.

28. Gray, C. J., Migas, L. G., Barran, P. E., Pagel, K., Seeberger, P. H., Eyers, C. E., Boons, G. J., Pohl, N. L. B., Compagnon, I., Widmalm, G., & Flitsch, S. L. (2019). Advancing Solutions to the Carbohydrate Sequencing Challenge. Journal of the American Chemical Society, 141(37), 14463–14479.

29. Grossutti, M., & Dutcher, J. R. (2016). Correlation between Chain Architecture and Hydration Water Structure in Polysaccharides. Biomacromolecules, 17(3), 1198–1204.

30. Guan, N., Blomsma, S. A., van Midwoud, P. M., Fahy, G. M., Groothuis, G. M. M., & de Graaf, I. A. M. (2012). Effects of cryoprotectant addition and washout methods on the viability of precision-cut liver slices. Cryobiology, 65(3), 179–187.

31. Guerreiro, B. M., Consiglio, A. N., Rubinsky, B., Powell-Palm, M. J., & Freitas, F. (2022). Enhanced Control over Ice Nucleation Stochasticity Using a Carbohydrate Polymer Cryoprotectant. ACS Biomaterials Science and Engineering.

32. Guerreiro, B. M., Freitas, F., Lima, J. C., Silva, J. C., Dionísio, M., & Reis, M. A. M. (2020). Demonstration of the cryoprotective properties of the fucose-containing polysaccharide FucoPol. Carbohydrate Polymers, 245, 116500.

33. Guerreiro, B. M., Silva, J. C., Lima, J. C., Reis, M. A. M., & Freitas, F. (2021). Antioxidant potential of the bio-based fucose-rich polysaccharide fucopol supports its use in oxidative stress-inducing systems. Polymers, 13(18), 3020.

34. Guerreiro, B. M., Silva, J. C., Torres, C. A. V., Alves, V. D., Lima, J. C., Reis, M. A. M., & Freitas, F. (2021). Development of a Cryoprotective Formula Based on the Fucose-Containing Polysaccharide FucoPol. ACS Applied Bio Materials, 4(6), 4800–4808.

35. Hietanen, J., & Salo, O. P. (1984). Binding of four lectins to normal human oral mucosa. European Journal of Oral Sciences, 92(5), 443–447.

36. Jiang, L., Kaoshan, C., Xuezheng, L., Peiqing, H., Guangyou, L., Jiang, L., Kaoshan, C., Xuezheng, L., Peiqing, H., & Guangyou, L. (2006). Production and characterization of an extracellular polysaccharide of antarctic marine bacteria Pseudoalteromonas sp. S-15-13. Acta Oceanologica Sinica, 2006, Issue 6, Pages: 106-115, 6, 106–115.

37. Kazak Sarilmiser, H., & Toksoy Oner, E. (2014). Investigation of anti-cancer activity of linear and aldehyde-activated levan from Halomonas smyrnensis AAD6T. Biochemical Engineering Journal, 92, 28–34.

38. Khudyakov, A. N., Polezhaeva, T. V., Zaitseva, O. O., Gunter, E. A., Solomina, O. N., Popeyko, O. V., Shubakov, A. A., & Vetoshkin, K. A. (2015). The Cryoprotectant Effect of Polysaccharides from Plants and Microalgae on Human White Blood Cells. Biopreservation and Biobanking, 13(4), 240–246.

39. Kratochvílová, I., Golan, M., Pomeisl, K., Richter, J., Sedláková, S., Šebera, J., Mičová, J., Falk, M., Falková, I., Řeha, D., Elliott, K. W., Varga, K., Follett, S. E., & Šimek, D. (2016). Theoretical and experimental study of the antifreeze protein AFP752, trehalose and dimethyl sulfoxide cryoprotection mechanism: correlation with cryopreserved cell viability. RSC Advances, 7(1), 352–360.

40. Li, B., Lu, F., Wei, X., & Zhao, R. (2008). Fucoidan: structure and bioactivity. *Molecules (Basel*, Switzerland), 13(8), 1671–1695.

41. Li, Y., Jiang, Y., Zhang, L., Qian, W., Hou, X., & Lin, R. (2021). Exogenous l-fucose protects the intestinal mucosal barrier depending on upregulation of FUT2-mediated fucosylation of intestinal epithelial cells. FASEB Journal, 35(7).

42. Limoli, D. H., Jones, C. J., & Wozniak, D. J. (2015). Bacterial Extracellular Polysaccharides in Biofilm Formation and Function. Microbiology Spectrum, 3(3).

43. Listinsky, J. J., Siegal, G. P., & Listinsky, C. M. (1998). Alpha-L-fucose: a potentially critical molecule in pathologic processes including neoplasia. American Journal of Clinical Pathology, 110(4), 425–440.

44. Liu, S. B., Chen, X. L., He, H. L., Zhang, X. Y., Xie, B. Bin, Yu, Y., Chen, B., Zhou, B. C., & Zhang, Y. Z. (2013). Structure and ecological roles of a novel exopolysaccharide from the Arctic sea ice bacterium Pseudoalteromonas sp. strain SM20310. Applied and Environmental Microbiology, 79(1), 224–230.

45. Ma, B., Simala-Grant, J. L., & Taylor, D. E. (2006). Fucosylation in prokaryotes and eukaryotes. Glycobiology, 16(12).

46. Martin-Pastor, M., Ferreira, A. S., Moppert, X., Nunes, C., Coimbra, M. A., Reis, R. L., Guezennec, J., & Novoa-Carballal, R. (2019). Structure, rheology, and copper-complexation of a hyaluronan-like exopolysaccharide from Vibrio. Carbohydrate Polymers, 222, 114999.

47. Matsumura, K., Rajan, R., & Ahmed, S. (2022). Bridging polymer chemistry and cryobiology. Polymer Journal 2022 55:2, 55(2), 105–115.

48. Mazur, P. (2010). A biologist’s view of the relevance of thermodynamics and physical chemistry to cryobiology. Cryobiology, 60(1), 4–10.

49. Murray, K. A., & Gibson, M. I. (2022). Chemical approaches to cryopreservation. Nature Reviews Chemistry 2022 6:8, 6(8), 579–593. 10.1038/s41570-022-00407-4

50. Nakamura, A., Takahashi, T., Yoshida, R., Maeda, H., & Corredig, M. (2004). Emulsifying properties of soybean soluble polysaccharide. Food Hydrocolloids, 18(5), 795–803.

51. Nwodo, U. U., Green, E., & Okoh, A. I. (2012). Bacterial exopolysaccharides: Functionality and prospects. International Journal of Molecular Sciences, 13(11), 14002–14015.

52. Péterszegi, G., Fodil-Bourahla, I., Robert, A. M., & Robert, L. (2003). Pharmacological properties of fucose. Applications in age-related modifications of connective tissues. Biomedicine & Pharmacotherapy, 57(5–6), 240–245.

53. Péterszegi, G., Isnard, N., Robert, A. M., & Robert, L. (2003). Studies on skin aging. Preparation and properties of fucose-rich oligo– and polysaccharides. Effect on fibroblast proliferation and survival. Biomedicine and Pharmacotherapy, 57(5–6), 187–194.

54. Péterszegi, G., Robert, A. M., & Robert, L. (2003). Protection by L-fucose and fucose-rich polysaccharides against ROS-produced cell death in presence of ascorbate. Biomedicine & Pharmacotherapy, 57(3–4), 130–133.

55. Qiu, Y., Hudait, A., & Molinero, V. (2019). How Size and Aggregation of Ice-Binding Proteins Control Their Ice Nucleation Efficiency. Journal of the American Chemical Society, 141(18), 7439–7452.

56. Rajan, R., Hayashi, F., Nagashima, T., & Matsumura, K. (2016). Toward a Molecular Understanding of the Mechanism of Cryopreservation by Polyampholytes: Cell Membrane Interactions and Hydrophobicity. Biomacromolecules, 17(5), 1882–1893.

57. Roeters, S. J., Golbek, T. W., Bregnhøj, M., Drace, T., Alamdari, S., Roseboom, W., Kramer, G., Šantl-Temkiv, T., Finster, K., Pfaendtner, J., Woutersen, S., Boesen, T., & Weidner, T. (2021). Ice-nucleating proteins are activated by low temperatures to control the structure of interfacial water. Nature Communications, 12(1).

58. Ruiz-Ruiz, C., Srivastava, G. K., Carranza, D., Mata, J. A., Llamas, I., Santamaría, M., Quesada, E., & Molina, I. J. (2011). An exopolysaccharide produced by the novel halophilic bacterium Halomonas stenophila strain B100 selectively induces apoptosis in human T leukaemia cells. Applied Microbiology and Biotechnology, 89(2), 345–355.

59. Russell, L., Waring, P., & Beaver, J. P. (1998). Increased cell surface exposure of fucose residues is a late event in apoptosis. Biochemical and Biophysical Research Communications, 250(2), 449–453.

60. Santarius, K. A., & Bauer, J. (1983). Cryopreservation of spinach chloroplast membranes by low-molecular-weight carbohydrates. I. Evidence for cryoprotection by a noncolligative-type mechanism. Cryobiology, 20(1), 83–89.

61. Schneider, M., Al-Shareffi, E., & Haltiwanger, R. S. (2017). Biological functions of fucose in mammals. Glycobiology, 27(7), 601.

62. Shtukenberg, A. G., Ward, M. D., & Kahr, B. (2017). Crystal Growth with Macromolecular Additives. Chemical Reviews, 117(24), 14042–14090.

63. Skjåk-Bræk, G., & Draget, K. I. (2012). Alginates: Properties and Applications. In Polymer Science: a Comprehensive Reference: Volume 1-10 (Vols. 1–10, pp. 213–220). Elsevier.

64. Stubbs, C., Bailey, T. L., Murray, K., & Gibson, M. I. (2020). Polyampholytes as Emerging Macromolecular Cryoprotectants. Biomacromolecules, 21(1), 7.

65. Stubbs, C., Lipecki, J., & Gibson, M. I. (2017). Regioregular Alternating Polyampholytes Have Enhanced Biomimetic Ice Recrystallization Activity Compared to Random Copolymers and the Role of Side Chain versus Main Chain Hydrophobicity. Biomacromolecules, 18(1), 295–302.

66. Sun, X., Wu, Y., Song, Z., & Chen, X. (2022). A review of natural polysaccharides for food cryoprotection: Ice crystals inhibition and cryo-stabilization. Bioactive Carbohydrates and Dietary Fibre, 27, 100291.

67. Sun, Y., Liu, J., Li, Z., Wang, J., & Huang, Y. (2021). Nonionic and Water-Soluble Poly(d / l – serine) as a Promising Biomedical Polymer for Cryopreservation. ACS Applied Materials and Interfaces, 13(16), 18454–18461.

68. Thomas, A. W., York, N., Stetten, D., Stetten, M. R., Bentley, O. G., & Kabat, E. A. (1956). Blood group substances: Their chemistry and immunochemistry. Journal of Chemical Education, 33(12), 653.

69. Thomson, L. K., Fleming, S. D., Aitken, R. J., De Iuliis, G. N., Zieschang, J. A., & Clark, A. M. (2009). Cryopreservation-induced human sperm DNA damage is predominantly mediated by oxidative stress rather than apoptosis. *Human Reproduction (Oxford*, England), 24(9), 2061–2070.

70. Torres, C. A. V., Ferreira, A. R. V., Freitas, F., Reis, M. A. M., Coelhoso, I., Sousa, I., & Alves, V. D. (2015). Rheological studies of the fucose-rich exopolysaccharide FucoPol. International Journal of Biological Macromolecules, 79, 611–617.

71. Trachtenberg, S. (1986). Conformation and aggregation of a polysaccharide: In solution, as transported in Golgi vesicles, and in an extracellular matrix. Journal of Ultrastructure and Molecular Structure Research, 97(1–3), 89–102.

72. Wang, C. S., Virgilio, N., Carreau, P. J., & Heuzey, M. C. (2021). Understanding the Effect of Conformational Rigidity on Rheological Behavior and Formation of Polysaccharide-Based Hybrid Hydrogels. Biomacromolecules, 22(9), 4016–4026.

73. Wang, H. M., Loganathan, D., & Linhardt, R. J. (1991). Determination of the pK(a) of glucuronic acid and the carboxy groups of heparin by 13C-nuclear-magnetic-resonance spectroscopy. Biochemical Journal, 278(3), 689–695.

74. Wang, H., Wang, X., & Wu, D. (2022). Recent Advances of Natural Polysaccharide-based Double-network Hydrogels for Tissue Repair. Chemistry – An Asian Journal, 17(17).

75. Wang, J., Salem, D. R., & Sani, R. K. (2019). Extremophilic exopolysaccharides: A review and new perspectives on engineering strategies and applications. Carbohydrate Polymers, 205, 8–26.

76. Xu, X. L., Li, S., Zhang, R., & Le, W. D. (2022). Neuroprotective effects of naturally sourced bioactive polysaccharides: an update. Neural Regeneration Research, 17(9), 1907.

77. Yang, J. S., Xie, Y. J., & He, W. (2011). Research progress on chemical modification of alginate: A review. Carbohydrate Polymers, 84(1), 33–39.

78. Ye, H., & Azar, D. (1998). Expression of gelatinases A and B, and TIMPs 1 and 2 during corneal wound healing. Investigative Ophthalmology & Visual Science.

79. Yeh, Y., & Feeney, R. E. (1996). Antifreeze proteins: Structures and mechanisms of function. Chemical Reviews, 96(2), 601–618.

80. Zeece, M. (2020). Food additives. Introduction to the Chemistry of Food, 251–311.

81. Zieske, J. D., & Bernstein, I. A. (1982). Modification of Cell Surface Glycoprotein: Addition Fucosyl Residues during Epidermal Differentiation of. The Journal of Cell Biology, 95, 626–631.

