## Supplementary Information for "Fucose is an essential feature in cryoprotective polysaccharides"

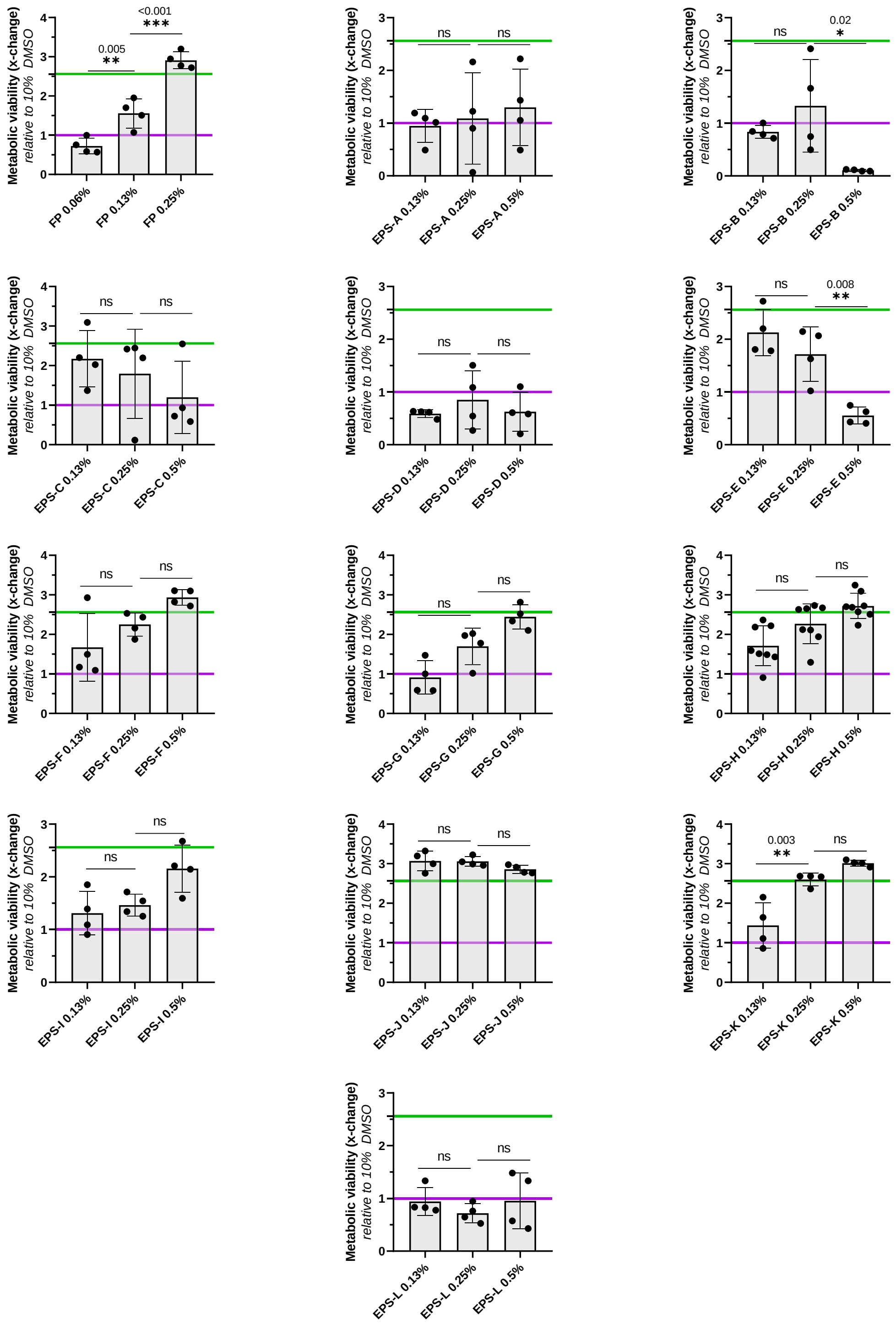

**Figure SI.1.** Post-thaw viability of Vero cells cryopreserved in a freezing medium containing the biotechnologically produced polysaccharides, at varying concentrations, supplemented with 10% DMSO. The purple and green thresholds represent 10% DMSO and CryoStor™ CS5 controls, respectively.

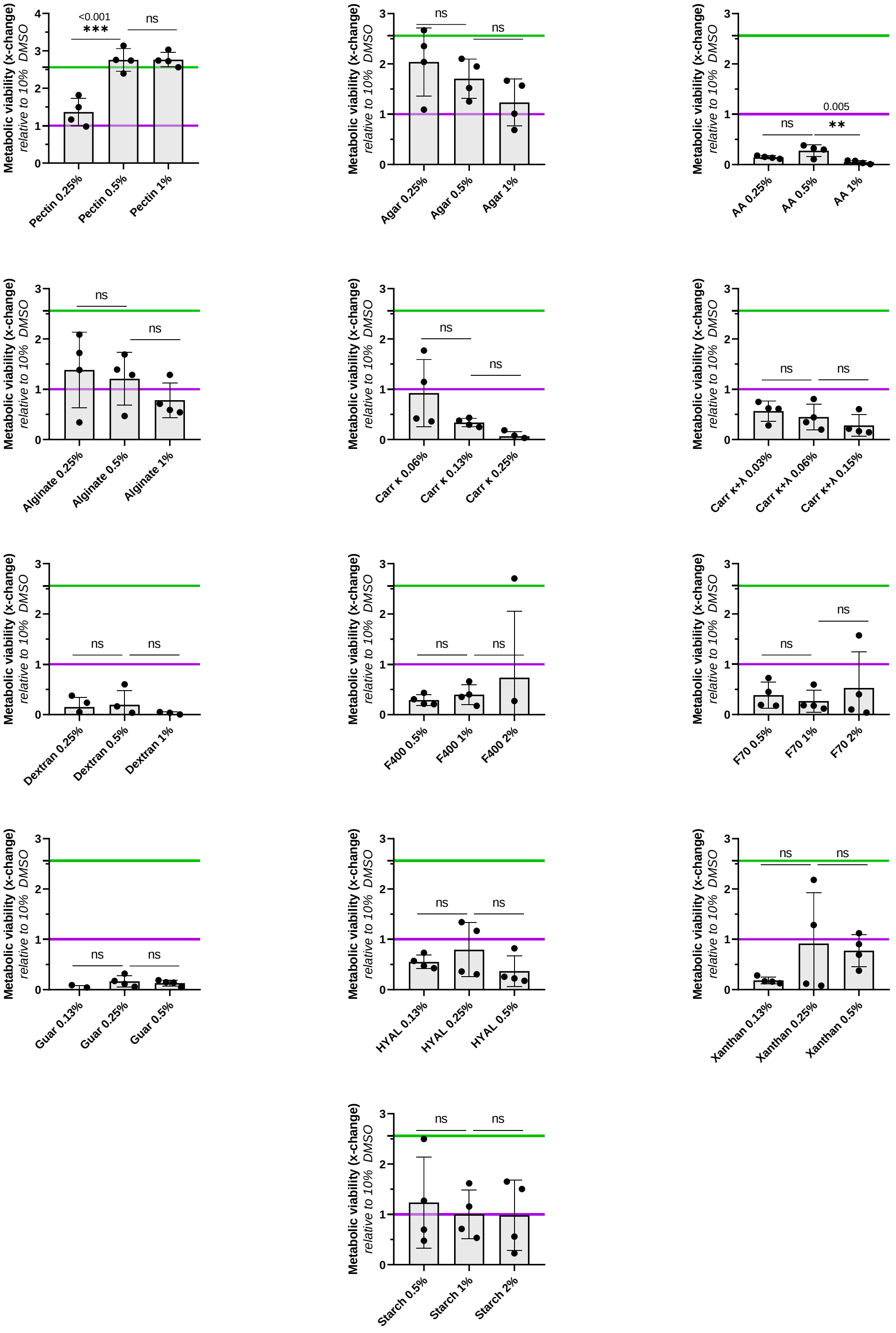

**Figure SI.2.** Post-thaw viability of Vero cells cryopreserved in a freezing medium containing the commercially sourced polysaccharides, at varying concentrations, supplemented with 10% DMSO. The purple and green thresholds represent 10% DMSO and CryoStor™ CS5 controls, respectively.

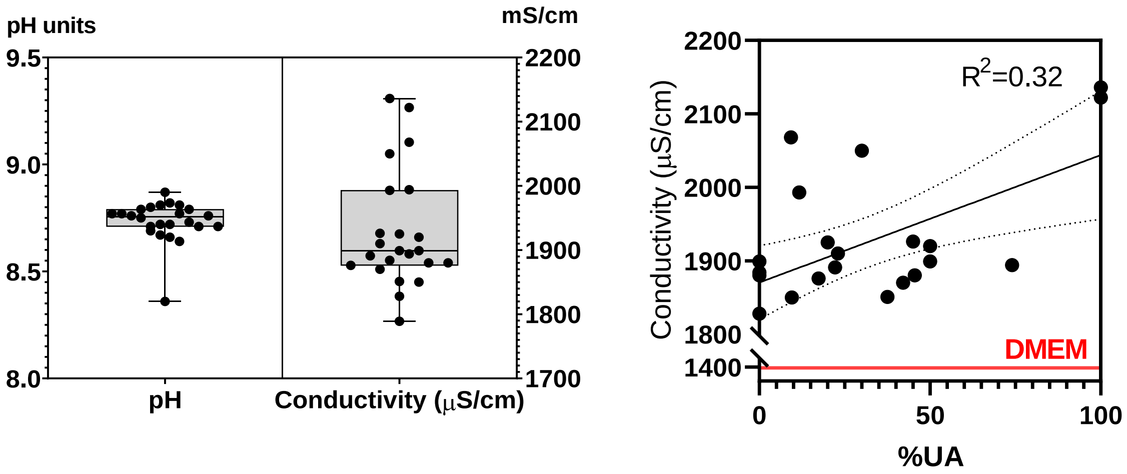


**Figure SI.3.** Physicochemical characteristics (pH, conductivity) of all polysaccharides studied, dissolved in DMEM culture medium. DMEM uses a sodium bicarbonate buffer system (3.7 g/L) to mitigate drastic pH changes. pH measurements were performed at room-temperature conditions, thus yielding average pH 8.8, but cells cultured in DMEM-based solutions are incubated in a 5% $CO_2$ atmosphere to maintain physiological pH, hence explaining the variation (*manufacturer information*).

### SI.2. Polysaccharide dataset description

**Table SI.1. Complete information on the polysaccharide dataset used in Vero cell cryopreservation experiments,** from which information on UA%, Fuc% and total Neutral% was gathered. Producing EPS strain designations, acyl and sulfate content, other infrequent residues and zero-shear viscosity values are shown.

| | Neutral monomers | | | | | Anionic monomers | | | | | | | | | | |
|---|---|---|---|---|---|---|---|---|---|---|---|---|---|---|---|---|
| EPS | **D-Glc** (mol%) | **D-Gal** (mol%) | **D-Man** (mol%) | **L-Fuc** (mol%) | **L-Rha** (mol%) | **D-GlcA** (mol%) | **D-GalA** (mol%) | **D-GulA** (mol%) | **D-ManA** (mol%) | **Other monomers** (mol%) | **Pyruvyl** (wt.%) | **Succinyl** (wt.%) | **Acetyl** (wt.%) | **Sulfate** (wt.%) | **MW** (MDa) | $\eta_0$ (Pa·s·%$^{-1}$) |
| ***Biotechnologically produced*** | | | | | | | | | | | | | | | | |
| **FucoPol** | 32.5±4.5 | 26±1 | - | 34±2 | - | 9.5±0.5 | - | - | - | - | 13.5±0.5 | 2.5±0.5 | 4.0±1.0 | - | 3.8±2.1 | 12 |
| **A** | 26.3 | 15.8 | 15.8 | - | - | 26.3 | 15.8 | - | - | - | 4.9±0.1 | - | 0.5±0.1 | 2.8±0.1 | 3.1±1.5 | 76.2 |
| **B** | 18.2 | 36.4 | - | - | - | 27.3 | 18.2 | - | - | - | 0.2 | 1.8±0.1 | 0.5 | 3.3±0.3 | 4.6 | 16.1 |
| **C** | 8.7 | 39.1 | 13 | 4.3 | 17.4 | 8.7 | 8.7 | - | - | - | 1.1±0.2 | - | - | 3.4 | 1.2 | 11.3 |
| **D** | 20 | 20 | 15 | - | - | 30 | 15 | - | - | - | - | - | 2.1 | 2 | 1.4 | 0.74 |
| **E** | 11.1 | 33.3 | 22.2 | - | 11.1 | 11.1 | 11.1 | - | - | - | - | 1 | - | 2.9 | 3.4±0.9 | 26.3 |
| **F** | 25 | 25 | - | - | 12.5 | 25 | 12.5 | - | - | - | 5.5±0.3 | - | 0.7 | 3.4 | 3.2 | 0.009 |
| **G** | $25^{-NAc}$ | $25^{-NAc}$ | - | - | - | 50 | - | - | - | - | - | - | - | - | 0.51 | - |
| **H** | 18 | 22 | - | 37 | - | 23 | - | - | - | - | 5.1 | 0.6 | 4.2 | - | 4.2 | - |
| **I** | 26.7±0.1 | 4.4±0.1 | 12.3±0.5 | - | - | - | - | - | - | GlcN (36.2±0.1), Rib (20.4±0.7) | 0.5±0.2 | - | 0.9 | 6.0±0.5 | 1.3±0.6 | - |
| J | 40.6±0.4 | 23.7±0.1 | - | 26.4±0.1 | - | 9.3±0.4 | - | - | - | - | - | - | - | - | 7.9 | - |
| K | 23.7±0.3 | 24.2±0.3 | - | 45.0±0.2 | - | 7.1±0.8 | - | - | - | - | - | - | - | - | - | - |
| L | 2.7 | 31.9 | 18.1 | 1.0 | - | 2.4 | 9.3 | - | - | Xyl, Ara (11, 22.8) | - | - | - | 10.6 | - | - |
| *Comercially sourced* | | | | | | | | | | | | | | | | |
| **Pectin** | - | - | - | - | - | - | 74†–100 | - | - | - | - | - | - | - | 0.18±0.1 | |
| **Agar** | - | 70† | - | - | - | 30† | - | - | - | - | likely | - | - | likely | 0.12 | |
| **G-alginate** | - | - | - | - | - | - | - | 75† | 25† | - | - | - | likely | - | 0.21±0.2 | |
| **M-alginate** | - | - | - | - | - | - | - | 35† | 65† | - | - | - | likely | - | 0.22 | |
| **Starch** | 100 | - | - | - | - | - | - | - | - | - | - | - | - | - | 0.3±0.2 | |
| **Carr. κ** | - | 100 | - | - | - | - | - | - | - | - | - | - | - | 19.8† | 0.25 | |
| **Carr. κ+λ** | - | 100 | - | - | - | - | - | - | - | - | - | - | - | 24.8† | 0.8† | |
| **Xanthan** | 40 | - | 40 | - | - | 20† | - | - | - | - | likely | - | - | - | 2 | |
| **Hyal. acid** | $50^{-NAc}$ | - | - | - | - | 50 | - | - | - | - | - | - | - | - | 0.85±0.4 | |
| **Ficoll®400** | 33 | - | - | - | - | - | - | - | - | Fru (33), epichlorohydrin (33) | - | - | - | - | 0.40† | |
| **Ficoll®70** | 33 | - | - | - | - | - | - | - | - | Fru (33), epichlorohydrin (33) | - | - | - | - | 0.07† | |
| **Dextran** | 100 | - | - | - | - | - | - | - | - | - | - | - | - | - | 0.04† | |
| **Guar** | - | 33 | 66 | - | - | - | - | - | - | - | - | - | - | - | 0.22 | |

† Specific manufacturer information. $^{-NAc}$ Monosaccharide residues N-acetylated.
